# The potential to breed for genetic legacy effects in sustainable farming systems

**DOI:** 10.1101/2025.11.03.686422

**Authors:** Shanice Van Haeften, Stephanie M. Brunner, Eric Dinglasan, Edward Fabreag, Joseph Eyre, Celine Mens, Ben J Hayes, Michael Udvardi, Samir Alahmad, Mitchell Eglinton, Ryan McQuinn, Merrill Ryan, Sarah van der Meer, Millicent R. Smith, Lee T. Hickey

**Affiliations:** Queensland Alliance for Agriculture and Food Innovation, The University of Queensland, QLD, Australia; School of Science, Western Sydney University, NSW, Australia; Queensland Department of Primary Industries, Warwick, QLD, Australia; School of Agriculture and Food Sustainability, Faculty of Science, The University of Queensland, QLD, Australia

## Abstract

Legume crops provide protein-rich food, critical disease breaks in cereal rotations, and contribute to soil fertility through symbiotic nitrogen fixation. However, crop improvement programs typically focus on within-crop performance rather than system-level benefits. We hypothesise that legacy effects (the influence of one crop’s genotype on subsequent crop performance) are under genetic control and could be leveraged in breeding programs. To test this, we evaluated how 309 genetically diverse mungbean genotypes influence subsequent wheat performance. The mungbean panel was grown, followed by a single wheat cultivar sown in the same plot locations. Remarkably, wheat yield varied by nearly 1 t ha□¹ (2.52-3.49 t ha□¹) depending solely on the preceding mungbean genotype, with legacy traits displaying moderate heritability (H²: 0.43-0.65), demonstrating untapped genetic potential for breeding. Analyses of mungbean traits, soil properties, and volatile organic compounds implicated root architecture, symbiotic nitrogen fixation and soil microbiome as potential biological drivers of legacy effects. Haplotype mapping identified genomic regions in mungbean associated with wheat yield and protein, revealing trade-offs between within-crop and legacy performance. Genetic simulations using empirically derived marker effects compared genomic selection strategies targeting mungbean yield, wheat yield, or both simultaneously. Balanced selection (50:50 weighting) achieved simultaneous gains in both crops (19.5% and 7.6%), highlighting the opportunity to breed for system-level productivity with reduced input requirements.

## Introduction

Modern crop breeding programs typically aim to optimise a single species, focusing on traits that enhance yield and quality, irrespective of the preceding or subsequent crops in rotation (Bennett et al., 2012). This approach achieved remarkable success during the Green Revolution, transforming wheat and rice production through high-yielding cultivars that fed billions globally, though often requiring increased fertiliser and irrigation inputs (Evenson and Gollin, 2003). Importantly, breeding remains effective even in low-input systems, as demonstrated by continued genetic gains across diverse production environments (Voss-Fels et al., 2019). However, the global population will approach 10 billion by 2050, and agricultural systems have become increasingly reliant on external inputs. Nitrogen fertiliser use has increased over 40% globally since 1990 (Li et al., 2025) and is projected to increase further to meet growing food demand, despite environmental and climate concerns. This calls for innovative breeding strategies to accelerate the development of more productive and sustainable crops (Hickey et al., 2019), particularly as the single-crop breeding paradigm may be missing critical genetic opportunities when the farming system is holistically considered.

Emerging evidence suggests that genetic variation within crop species creates measurable effects on subsequent crops in rotation (Rose et al., 2010; Ellouze et al., 2013), yet the genetic basis of these “legacy effects” remains largely unexplored. Legacy effects occur when the genetic characteristics of one crop influence the performance of subsequent crops through persistent changes to soil conditions that carry over between growing seasons (Somenahally et al., 2018; Lilley and Kirkegaard, 2016). More specifically, they are hypothesised to be genetically mediated via plant-soil feedback mechanisms involving water and nutrient dynamics, and microbiome assembly and function. Both the preceding crop (donor) and the subsequent crop (acceptor) genomes shape outcomes, where donor genes modify soil processes, such as through symbiotic nitrogen fixation, that interact with acceptor genes, such as N transporters, across seasons. The genomic basis of a legacy effect therefore refers to heritable genetic variation in the donor crop that drives differences in legacy outcomes, regardless of the specific agronomic pathway through which that influence operates.

While the benefits of crop rotations at the cross-species level are well established, for example legumes fixing nitrogen for cereals (e.g. Zhao et al., 2022), crop rotations to break disease cycles (e.g. Kirkegaard et al., 1997), and diverse root systems improving soil structure (e.g. Yu et al., 2025), the genetic variation within species that could enhance these rotational benefits through targeted breeding remains untapped in current breeding programs. Lilley and Kirkegaard (2016) highlight this concept, demonstrating that wheat cultivars with deeper, more extensive root systems extracted additional subsoil water but left soils drier at subsequent crop sowing, reducing predicted yield benefits when legacy effects were considered. Thus, selection for deeper root architecture within wheat breeding germplasm could lead to higher yielding wheat cultivars, but they may result in unintended negative consequences for subsequent crops. Integrating legumes into crop rotations enriches soil nitrogen through symbiotic nitrogen fixation, benefiting subsequent crops with yield increases often exceeding 10% and with substantial reductions in fertiliser requirements (Cernay et al., 2018; Zhao et al., 2022). However, the extent to which genetic variation within legume crop species modulates both positive and negative legacy effects remains unexplored.

We hypothesise that the genetic architecture of a crop can directly modulate the performance of the subsequent crop, creating legacy effects that can be leveraged through breeding to optimise integrated farming system productivity, ideally both as reduced inputs and increased yield across both crops (Fig 1). To test this, we conducted a proof-of-concept study using mungbean (*Vigna radiata*) as a preceding legume crop in a wheat-based rotation system. Mungbean contributes to the system by fixing nitrogen and providing a disease break, thereby enhancing the productivity and health of the subsequent wheat crop (Ilyas et al., 2018; Foyer, Nguyen & Lam, 2019). We evaluated how 309 genetically diverse mungbean accessions influenced performance of a subsequent wheat crop rotation, combined with haplotype mapping to identify genomic regions influencing both mungbean traits and legacy effects on wheat productivity and quality. Through genetic simulations using real haplotype effects, we further evaluated the potential of breeding programs that systematically consider traits which infer legacy effects (also known as ‘legacy traits’) alongside traditional performance metrics. These simulations were used to determine whether such integrated approaches could improve genetic gain and productivity for both crops through enhanced resource-use efficiency.

**Fig 1.**
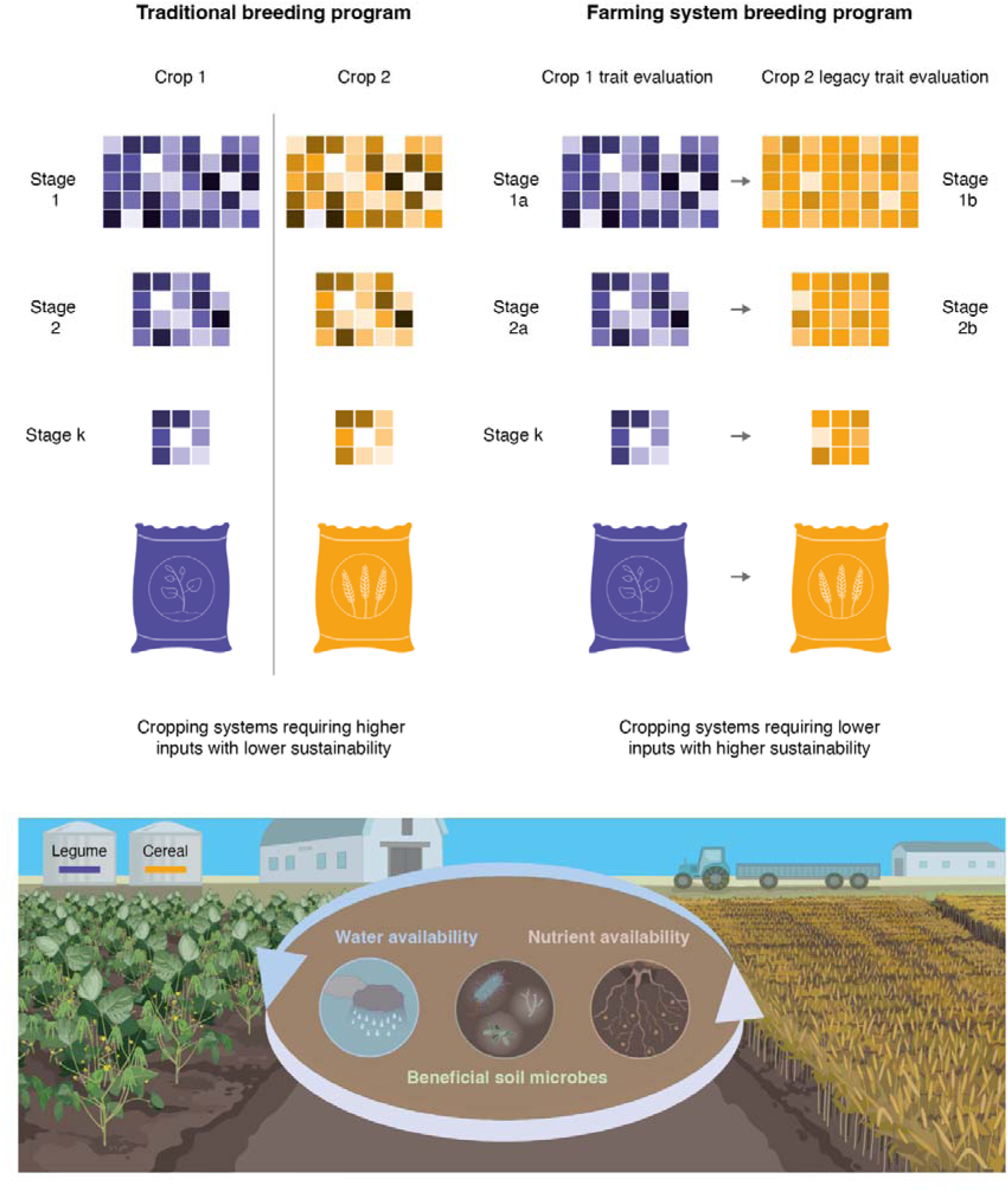
Conceptual framework for integrating legacy traits into farming systems breeding. Comparison of traditional breeding (optimising crop performance in isolation) with farming systems breeding that considers genetic effect of one crop on the next (legacy traits) within rotational systems. Different shades of dark purple to white represent genetic variation within Crop 1, whereas different shades of dark brown to white represent the subsequent (legacy effect) on Crop 2. In the farming systems approach, Crop 1 is evaluated for both within-crop performance traits and legacy effects on Crop 2 (Stage 1a), followed by assessment of Crop 2’s legacy trait expression (Stage 1b), with dual evaluation continuing through subsequent breeding stages. Providing a suggestion for breeding crops simultaneously to ensure favourable legacy traits increasing yield and improving sustainability. The lower panel illustrates how considering legacy traits could enable selection of genotypes that enhance system productivity through improved water-use efficiency, nutrient-use efficiency, and beneficial soil properties, including microbiome. This framework represents a potential paradigm shift toward multi-crop, multi-season breeding supporting sustainable intensification with reduced input requirements due to breeding for systems which improve or maintain the soil system for the following crop

## Results

### Mungbean genotype drives substantial heritable variation in wheat legacy performance

Wheat yield varied substantially by preceding mungbean genotype (2.52 to 3.49 t ha□¹; Crystal 3.06 t ha□¹), with individual mungbean genotypes enhancing subsequent wheat yield by up to 45% or reducing it by as much as 50% relative to Crystal (Fig. 2b). The mungbean genotype that produced the highest wheat yield was AGG325661 (3.49 ± 0.16), in comparison with mungbean genotype AGG325603 that produce the lowest wheat yield (2.52 ± 0.15). Wheat legacy traits showed moderate heritability (H² = 0.43–0.65), indicating that a substantial portion of variation in wheat performance was attributable to mungbean genotype. Wheat protein content and wheat NDRE at flowering also varied across mungbean genotypes (Supplementary Table 1). Environmental conditions during both the mungbean and wheat growing seasons were typical for the region with no major weather events (Supplementary Figure 1).

**Fig 2.**
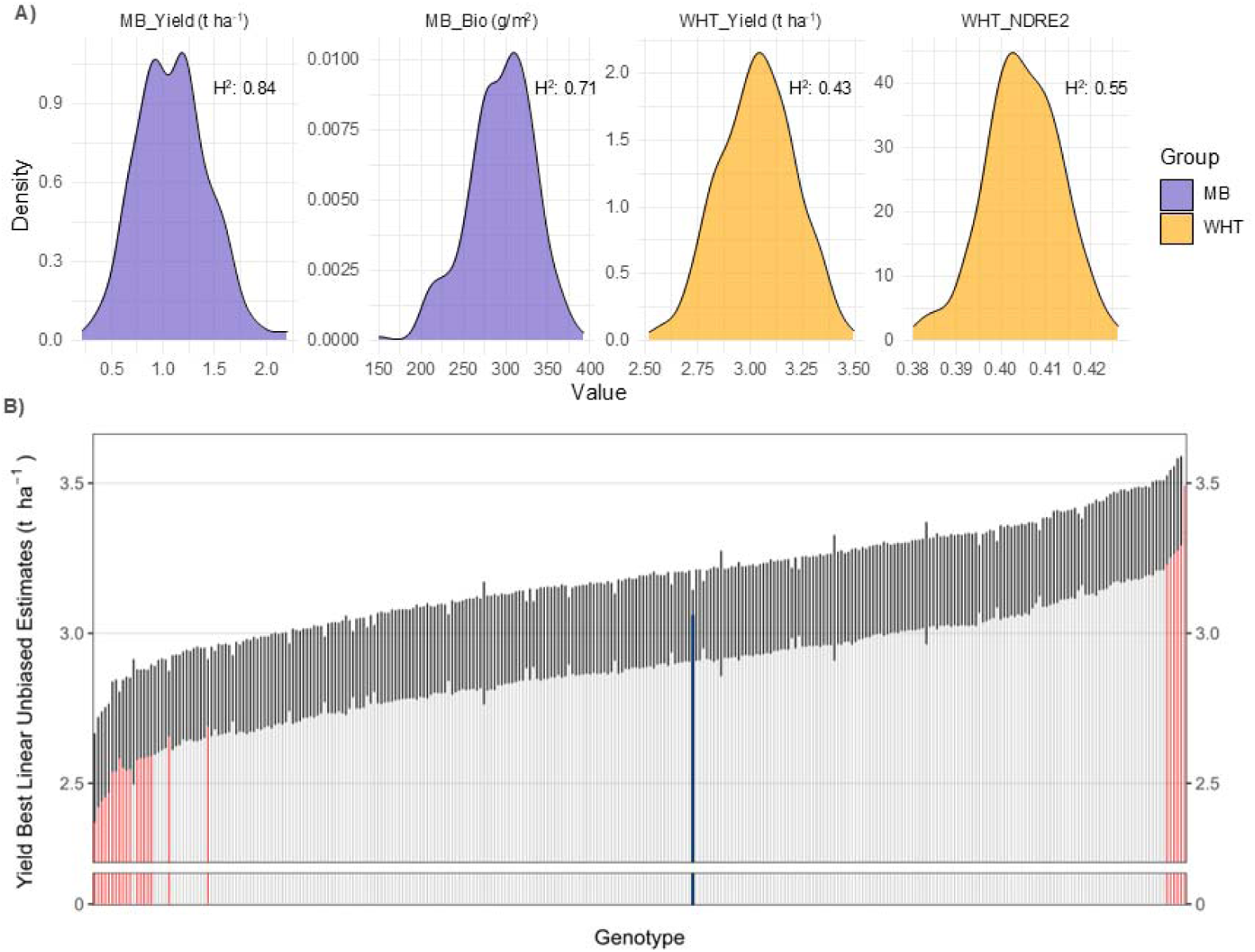
Genotypic variation and heritability of mungbean and wheat legacy traits. a) Distribution of phenotypic values for mungbean traits (yield and predicted biomass at flowering; purple) and wheat legacy traits (yield and NDRE at flowering; orange) across 309 genotypes. Broad-sense heritability (H²) for each trait is displayed in the top-right corner. b) Wheat yield performance ordered by increasing yield (t ha^-1^), reflecting legacy effects from preceding mungbean genotypes. Each bar represents the BLUE ± standard error for wheat following an individual mungbean genotype. Blue bar indicates wheat following the check cultivar ‘Crystal’; red bars indicate wheat yields significantly different from ‘Crystal’ (p < 0.05).

To explore mungbean traits potentially driving this legacy effect variation, we characterised phenotypic variation across the mungbean mini-core panel (Fig. 2a; Supplementary Tables 1 and 2). Mungbean yield varied 11-fold from 0.21 to 2.19 t ha□¹ (Crystal 1.31 t ha□¹) and aboveground biomass at mid canopy development showed 2.6-fold variation (151.1 to 393.3 g m□²; Crystal 280.7 g m□²). Flowering time ranged from 41 - 57 DAS across genotypes; Crystal 41 DAS (Supplementary Table 1). Mungbean NDRE at 90% black pod also showed considerable genotypic diversity across the panel (Supplementary Table 1). All mungbean within-crop traits exhibited high heritability (H² = 0.63–0.84), indicating that the variation observed reflects genuine differences among genotypes.

### Interactions between within-crop and legacy traits reveal trade-offs

Relationships between mungbean and wheat legacy traits varied in directionality and strength when evaluated across phenotypic, genetic, and haplotype levels (Fig. 3). At the phenotypic level (Fig 3a and 3b), mungbean yield showed a moderate negative correlation with wheat yield (*r* = −0.4, p < 0.05), suggesting potential trade-offs between mungbean productivity and subsequent crop performance. Wheat NDRE at flowering and wheat yield were positively correlated (*r* = 0.75, p < 0.05), indicating consistency among wheat physiological performance metrics, while wheat NDRE at flowering showed a significant negative correlation with mungbean yield (*r* = −0.48, p < 0.05).

**Fig 3.**
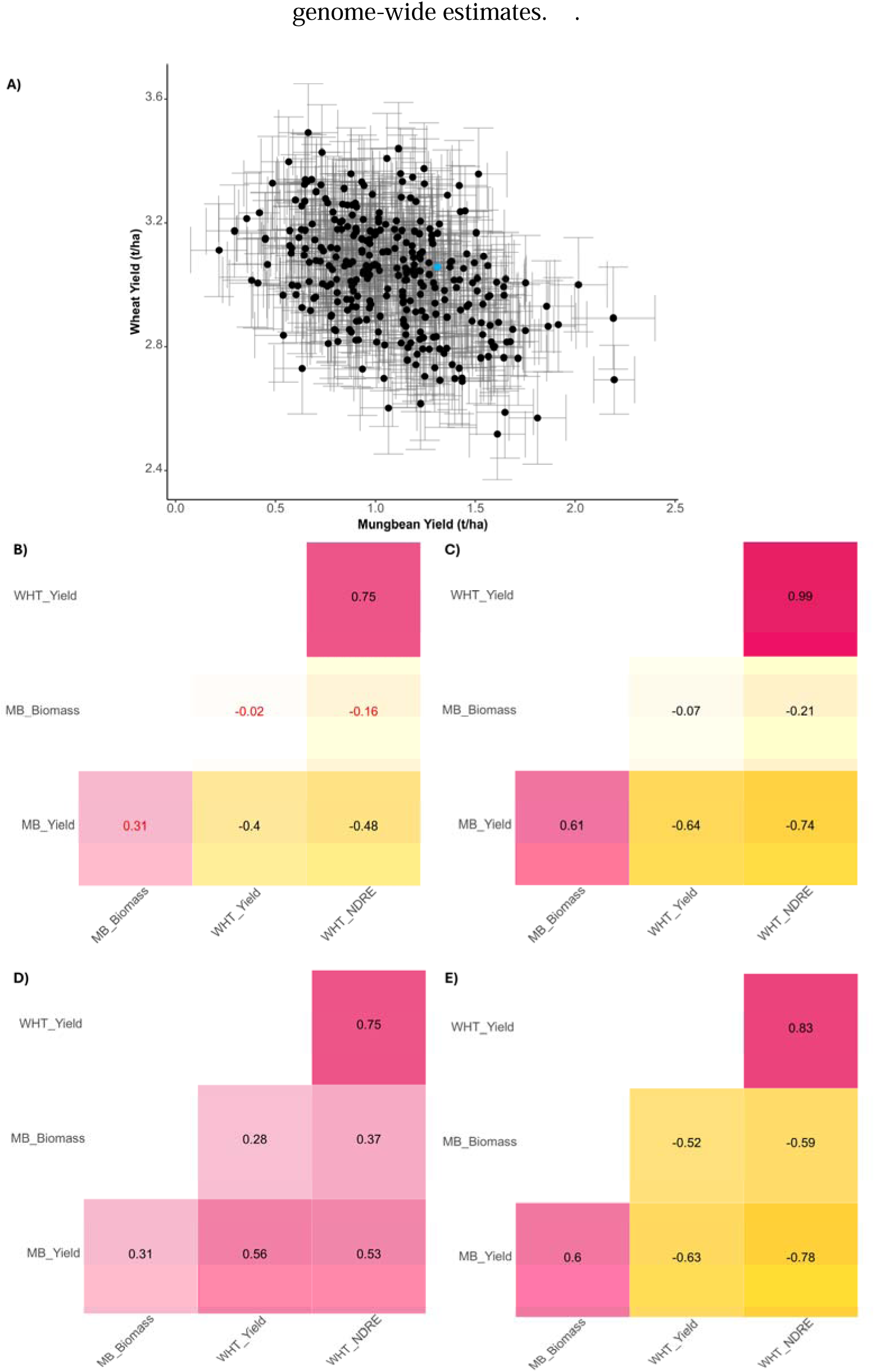
Correlations among mungbean traits and wheat legacy traits at phenotypic, genetic, and haplotype levels. Panels progressively remove environmental noise and complex interactions to reveal the genetic relationships between traits in the simplest form. A) Mungbean Yield BLUEs and Wheat Yield BLUEs with standard error and key variety Crystal highlighted in blue. B) Phenotypic correlations (r) based on BLUEs across all genotypes. C) Genetic correlations estimated from mixed-model variance components, indicating shared genetic architecture between traits. D) Haplotype-level variance correlations, indicating shared genomic regions contributing to trait variation. E) Haplotype effect correlations, quantifying concordance in direction and magnitude of haplotype influences across traits. Colour scale ranges from 1 (pink, positive correlation) through -1 (yellow, negative correlation). Non-significant correlations (p > 0.05) are indicated in red text.

Genetic correlations revealed stronger associations, highlighting underlying genetic relationships not evident at the phenotypic level (Fig 3b). Mungbean yield showed a strong positive genetic correlation with mungbean biomass (4 = 0.61) and a weak to moderate negative genetic correlation with wheat yield r = -0.29) and wheat NDRE (r = -0.21), suggesting that the genetic architecture underlying mungbean yield is mildly antagonistic to wheat performance.

Haplotype-level variance correlations were uniformly positive across all mungbean and wheat traits (*r* = 0.28–0.75, p < 0.05), indicating that shared genomic regions contribute to trait variation across the mungbean–wheat system (Fig 3c). Haplotype effect correlations revealed more nuanced patterns, with directional shifts between traits consistent with the genetic correlations observed (Fig 3d). Mungbean yield showed strong negative haplotype effect correlations with wheat yield (*r* = −0.63, p < 0.05) and wheat NDRE (*r* = −0.78, p < 0.05), but a positive association with mungbean biomass (*r* = 0.60, p < 0.05). These contrasting directions highlight key haplotypes that differentially influence mungbean performance and legacy benefits to wheat. Haplotype effect correlations between mungbean yield and wheat traits were considerably stronger than the corresponding genome-wide genetic correlations, reflecting the polygenic nature of these traits and the capacity of haplotype-level analysis to resolve region-specific genetic antagonism that is obscured in genome-wide estimates.

We selected the top 1% of haploblocks with highest variance for each of the six traits (12 per trait), which identified 33 unique haploblocks across all traits (Supplementary Table 3). Of these, 17 haploblocks were unique to single traits, with eight showing high variance for wheat legacy traits but not for any measured mungbean traits (Fig 4a; Supplementary Fig 2). The remaining 16 haploblocks showed high variance for at least two traits across both wheat and mungbean, with one haploblock (b000055, chr 1) showing high variance for all six traits, and three additional haploblocks, (b000160 chr 2; b000221 chr 3; and b000509 chr 5), showing high variance for five of six traits, suggesting these genomic regions harbour loci with pleiotropic effects within mungbean, and genetically mediated legacy effects influencing wheat traits in rotation systems. Among the top haploblocks, five were shared between mungbean yield and wheat yield (Fig. 4a): b000022 (chr 1), b000055 (chr 1), b000221 (chr 2), b000253 (chr 2), and b000509 (chr 5), while seven high-variance haploblocks did not overlap between crops. This pattern indicates both shared and independent genetic control of within-crop versus legacy traits.

**Figure 4.**
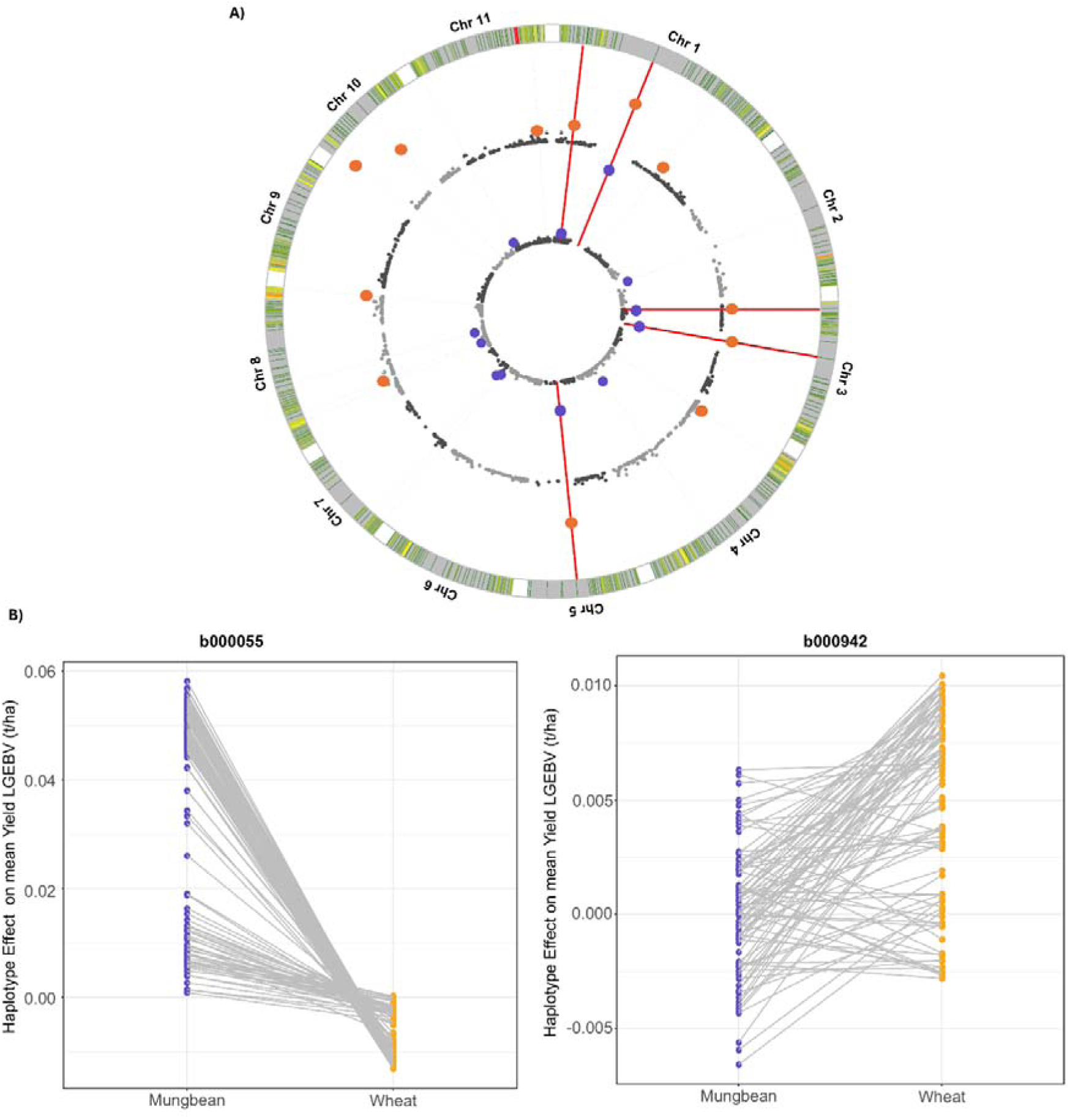
Haploblocks associated with mungbean yield and legacy effects on wheat yield. a) Circos plot displaying haploblock effect variance for yield in mungbean (inner ring) and wheat (outer ring) across the 11 mungbean chromosomes. The top 1% of haploblocks by variance are highlighted for each crop (mungbean purple, wheat orange). Red lines connect shared high variance haploblocks between crops, indicating loci with pleiotropic effects on system-level productivity. b) Haplotype effects for mungbean and wheat yield at two representative haploblocks: b000055 (chr 1) showing trade-offs, and b000942 (chr 10) showing aligned effects across both species. Points represent the unique haplotypes within the haploblock present in the population with lines connecting the same haplotype to visualise its effect on mungbean and wheat respectively.

Functional genes associated with nitrogen fixation in legumes were positioned within five top haploblocks identified in this study (Supplementary Table 4). Co-located genes were associated with broader root development processes, including gibberellin signalling (*MtDELLA2*), root meristem maintenance (*MtPLT1–4*), and auxin transport (*MtPIN2–4*), suggesting that root architecture may be a key mechanism underlying legacy effects.

To dissect these relationships at higher resolution, we examined two haploblocks in detail: b000055 (chr 1), the only haploblock with high variance for all six traits, and b000942 (chr 10), a haploblock with high variance for wheat yield but not ranking among the top 12 for mungbean yield. At haploblock b000055, haplotypes with the strongest positive effects for mungbean yield were associated with greater negative effects on wheat yield, consistent with the overall negative genetic correlation (Fig. 4b). This pattern illustrates the genomic basis of trade-offs between within-crop performance and legacy benefits. In contrast, block b000942 was not among the highest-variance regions for mungbean yield but nonetheless exhibited haplotype variance comparable to that observed for wheat, where it ranked among the top variance haploblocks. This haploblock harboured haplotypes with positive effects for both mungbean and wheat yield, as well as variants with neutral effects, indicating opportunities to select haplotypes beneficial for system-wide productivity without compromising individual crop performance.

### Simulated selection strategies reveal challenges and opportunities for system-level yield optimisation

To evaluate the potential for simultaneous genetic improvement of mungbean and its legacy effects on wheat, we simulated genomic selection strategies using variable weighting schemes for the two crops over 25 breeding cycles, using the real haplotype effects as a starting point. Simulated progenies were generated by sampling chromosomes with recombination from selected parents (e.g. Villiers et al., 2022). When haplotypes were selected solely to maximise mungbean yield (100% mungbean weighting), predicted genomic estimated breeding values for mungbean increased by 36.86% over the 25 cycles, with a corresponding 7.71% reduction in wheat yield, highlighting the presence of antagonistic haplotypes that benefit mungbean but negatively impact wheat performance.

Adjusting the selection index to include moderate wheat emphasis (70:30 mungbean:wheat weighting) maintained high genetic gain in mungbean (36.32%) while moderating the negative legacy impact on wheat (reduced to 4.59% loss), demonstrating a more balanced trade-off between within-crop and legacy benefits. In contrast, placing greater emphasis on wheat legacy performance (30:70 weighting) yielded strong improvements in wheat legacy yield (13.44% increase) with a modest decline in mungbean yield (22.56% reduction), indicating that prioritising legacy traits can substantially benefit the subsequent crop, albeit at a considerable cost to the initial crop.

Selection based on equal weighting (50:50) resulted in simultaneous yield gains for both mungbean and wheat (19.45% and 7.55%, respectively), though the overall magnitude of improvement was lower than in crop-specific optimisation scenarios. This balanced approach demonstrates the potential of system-wide genetic improvement through integrated breeding strategies that consider both within-crop performance and legacy effects. The simulation results reveal that while trade-offs exist between optimising individual crop performance versus rotational benefits, strategic selection can achieve meaningful gains across the entire cropping system, providing a pathway for sustainable intensification through breeding.

**Fig 5.**
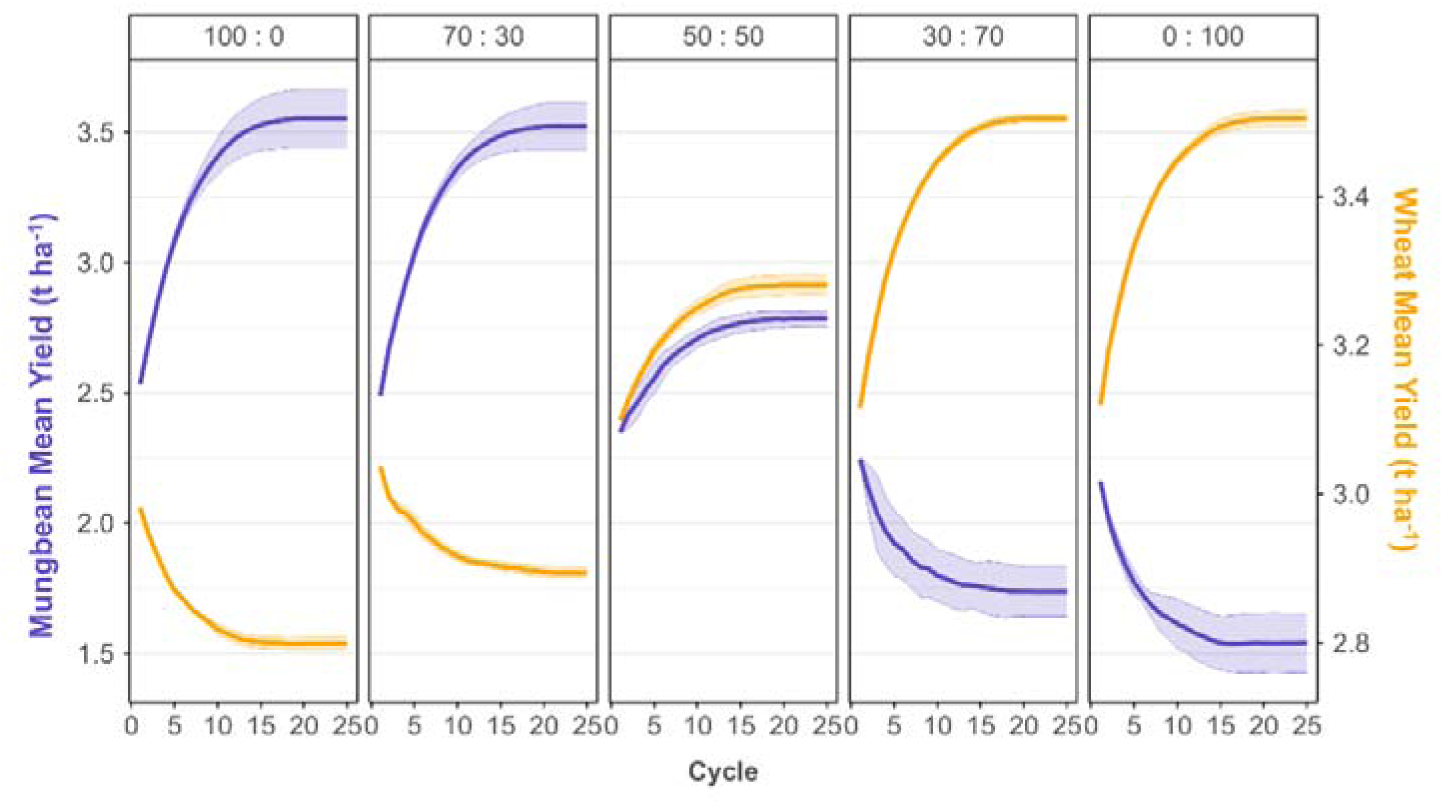
Simulated selection strategies reveal trade-offs and optimisation potential for breeding mungbean with system-level benefits. Genomic estimated breeding values (GEBVs) for mungbean yield (purple) and wheat legacy yield (orange) over 25 breeding cycles under five selection strategies. From left to right: 100% mungbean yield, 70:30 mungbean:wheat, 50:50 equal weighting, 30:70 mungbean:wheat, and 100% wheat legacy yield. Lines show mean GEBVs with 95% confidence intervals (shaded regions).

### Analysis of soil properties reveals genetic variation for biological drivers of legacy effects

To further explore traits potentially contributing to legacy effects, we sampled and analysed soil from a subset of 10 genetically diverse mungbean genotypes after mungbean harvest and prior to wheat sowing. Among the soil nutrient parameters assessed, nitrate (NO□□) levels differed significantly between genotypes (*p* = 0.01), indicating genotype-dependent variation in soil nitrogen dynamics. In contrast, no significant genotypic differences were detected for other soil nutrient components, including ammonium, phosphorus, or exchangeable cations (Supplementary Table 5).

To investigate possible differences in soil microbiome associated with mungbean genotype, we analysed soil volatile organic compounds (VOCs) for the same subset (Supplementary Table 6). Out of 143 VOCs detected in soil samples, 25 passed quality filter criteria, of which eleven showed significant genotypic effects (Supplementary Fig. 3). Dehydrocamphor and 2-methylisoborneol displayed the largest genotypic differences, with cultivar Crystal showing elevated abundance compared to bare soil control and other genotypes (Supplementary Fig. 4 and Supplementary Table 7). Berken displayed a distinct VOC profile with elevated levels of multiple compounds including 1,4-cis-dimethylcyclooctane, 4,5-dimethylnonane, and 3,7-dimethyloctan-1-ol compared to other genotypes and bare soil. Several mungbean genotypes were associated with significantly reduced accumulation of specific VOCs (3-methoxy-3-methylbutanol and styrene) compared to bare soil (Supplementary Fig. 4 and Supplementary Table 7), indicating potential for genotypic regulation of soil microbiome diversity.

## Discussion

While agricultural research has long demonstrated the value of crop rotations, this study reveals for the first time the potential for rotational benefits to be enhanced through targeted plant breeding by selecting genotypes that modulate the soil environment in ways that influence the performance of subsequent crops. To date, research has focused on species-level benefits (e.g., Li et al., 2019; Lago-Olveira et al., 2023; Salvagiotti et al., 2024), such as nitrogen fixation or disease suppression, rather than trait variation within-crop that influences legacy effects. Our results show that crop legacy effects are under genetic control.

In our mungbean-wheat rotation experiment, substantial phenotypic variation in wheat NDRE and yield, and to a lesser extent protein content, was observed as a function of the preceding mungbean genotype. Importantly, these legacy traits exhibited moderate heritability (H² = 0.43–0.65), demonstrating that they are amenable to selection. Remarkably, mungbean genotypes enhanced subsequent wheat yield by up to 45% or reduced it by 50% relative to the commercial check variety, revealing substantial genetic variation in legacy effects beyond environmental variation alone. These results highlight a real limitation in traditional breeding that focuses on maximising the performance within-crop, making an implicit assumption that any effect on subsequent crops is negligible. In contrast, our results demonstrate that legacy effects vary across genotypes and can have a substantial influence on subsequent crop performance. This reveals an opportunity for a fundamental shift in breeding philosophy, from exploiting fixed species-level rotational benefits to harnessing genetic variation within species for enhanced system-wide productivity.

It is important to emphasise that these heritability estimates are derived from a a single environment. While mungbean genotypes captured a proportion of the variance in wheat performance, these estimates are specific to the conditions of this single experiment. Environmental factors such as rainfall, temperature and soil moisture can interact with crop growth and development and may substantially modify the expression of legacy traits which in turn can affect the legacy effects in the following crop. For example, climate gradients across locations influence soil microbiome populations, with communities collected from warm and cool sites showing different functional effects on bud break phenology in trees (Ware et al., 2021). Consequently, genotype by environment interactions for both mungbean and wheat are likely to play an important role in determining the stability of legacy effects across production systems. Multi-environment trials represent an important next step to quantify the robustness of these genetic effects across varying climatic and soil conditions, now that the heritable nature of legacy effects has been determined.

This study also demonstrates that legacy effects can be genetically mapped to the genome of the preceding crop. The legacy traits investigated in this case study exhibited complex genetic architecture, with many agronomic traits showing shared genetic control. We identified both shared and unique haploblocks influencing mungbean traits and legacy effects on wheat, with 16 high-variance haploblocks showing overlapping effects across both crops. The magnitude of haplotype effects was consistently greater for mungbean traits than for legacy impacts on wheat. This is somewhat expected, as there are many other factors contributing to wheat performance beyond legacy effects, including the underlying genetics of the wheat cultivar and seasonal growing conditions. Several top haploblocks contained functional genes associated with root development processes, including gibberellin signalling (*MtDELLA2*), root meristem maintenance (*MtPLT1–4*), and auxin transport (*MtPIN2–4*), rather than nodulation-specific pathways. This suggests that some of the legacy traits identified in this study may be mediated through root growth and architecture, which influence soil nutrient and water status and rhizosphere interactions, providing a potential mechanistic link between mungbean genotype and subsequent wheat performance. It is important to note that many of these haploblocks are large and likely contain multiple functional genes across many pathways; the identification of root-specific candidates therefore provides supporting evidence for a root-mediated mechanism, but molecular validation will be required to ascertain the specific genes involved.

Trade-offs between mungbean yield and legacy effects on wheat performance indicate that traits conferring high productivity in one crop may compromise the next crop in rotation. Specifically, high-yielding mungbean genotypes were associated with reduced wheat NDRE at flowering, which in turn was linked to lower wheat yield, consistent with a source-sink limitation driven by environmental modification. This suggests that mungbean genotypes may alter the soil environment through mechanisms such as differential nutrient extraction, altered water availability, or rhizosphere-mediated shifts in the soil microbiome. These changes may reduce resource availability for the subsequent crop, thereby impacting biomass accumulation and final yield. Nitrogen availability and uptake are potential sources of the observed trade-offs though the N uptake partitioning characteristics of different mungbean cultivars are not well explored. Additional experiments could address the N partitioning of accessions and the correlation with legacy effects for wheat.

Genomic analysis revealed negative correlations between mungbean haplotype effects on mungbean and wheat traits, with particularly strong trade-offs for yield. For instance, haploblock b000055 showed that haplotypes associated with the highest mungbean yield also had the most detrimental effect on legacy wheat yield, whereas b000942 included haplotypes with neutral or even positive effects on both crops. Importantly, these ‘correlation breakers’ occurred mostly in the lower effect size range, suggesting that system-level improvement may be best achieved through genome-wide strategies like weighted genomic selection or haplotype stacking, rather than marker-assisted selection for large-effect loci only.

Although the precise biological drivers of these legacy effects remain unclear, our targeted soil analyses indicate that there is measurable variation in both nutrient availability and VOC profiles among diverse mungbean genotypes (Supplementary Tables 5 and 7). While the concept of microbes utilising VOCs for interkingdom communication is relatively recent (Weisskopf, et al., 2021), soil volatilomic analysis holds promise as a cost-effective means to assess soil health and microbial biodiversity. Several genotype-associated VOCs were identified, including 2-pentylfuran, which is known to be emitted from crop roots and acts as an insect attractant (la Forgia, et al., 2023), and 2-methylisoborneol and related terpenes, which may indicate the presence of specific bacterial types such as actinobacteria, myxobacteria, and cyanobacteria (Weisskopf et al., 2021; Martin-Sanchez, et al., 2019). While directly linking these VOCs to specific microbes or plant processes requires additional investigation, these findings highlight the potential for genotypic influence on the soil environment through root architecture, nutrient and water acquisition, or microbial community structuring. In particular, legacy effects mediated through persistent changes in the rhizosphere microbiome are a promising avenue, as legume genotypes are known to establish distinct microbial communities that can enhance nutrient cycling and disease suppression for subsequent crops (Yu et al., 2025; Kirkegaard et al., 1997). VOC profiles are used here as indicators of potential genotype-dependent differences in soil microbial activity rather than as direct evidence of specific microbial processes or community composition. Future work employing targeted microbiome characterisation approaches, such as shotgun metagenomics, will be necessary to resolve the specific microbial taxa and functional pathways underlying these genotype-dependent soil legacies. Together, these findings underscore the need for further biological characterisation of legacy mechanisms and demonstrate the feasibility of incorporating legacy traits into breeding strategies to optimise productivity across crop rotations.

The capacity of plants to modify their soil environment through root architecture and rhizosphere interactions may have deep evolutionary roots. Recent work in model systems demonstrates that natural genetic variation in plant species can shape soil microbial communities involved in nitrogen cycling, suggesting that niche construction through soil modification is a heritable, selectable trait (Przybylska et al., 2024). Domestication processes that prioritised yield may have inadvertently eroded this capacity by reducing selection pressure on traits mediating plant-soil feedbacks. The substantial genetic variation for legacy effects observed here across a diverse mungbean panel, including landraces and diverse accessions, is consistent with this interpretation, and suggests that beneficial alleles for legacy effect capacity may be recoverable through strategic use of pre-breeding diversity. Against this backdrop, simulated genomic selection strategies reveal the practical consequences of ignoring legacy effects in breeding.

When selection was fully weighted toward mungbean yield, substantial genetic gain in mungbean (36.86%) was observed but consistent reductions in wheat legacy yield (7.71%) were noted, reflecting negative genetic correlations between traits expressed in the two crops. In contrast, selection indices designed to balance genetic gain in both crops moderated these trade-offs (70:30 weighting) or eliminated them (50:50 weighting). While absolute genetic gain in each crop was reduced, overall system performance improved by minimising negative legacy effects. Equal weighting (50:50) achieved simultaneous yield gains for both mungbean and wheat (19.45% and 7.55%, respectively), demonstrating that within-crop and legacy traits under negative correlation can be successfully managed through weighted selection indices. In practice, optimal weighting schemes could consider economic factors including relative commodity prices, input costs such as nitrogen fertiliser, and farm-specific production costs, as the economic value of legacy effects may vary substantially depending on market conditions and production systems (Melton et al., 1979). These simulations provide a framework for integrating legacy traits into breeding pipelines, offering insights into how genomic prediction can be optimised to support multi-season, multi-crop production.

The implications of breeding for legacy traits extend far beyond the immediate case study, presenting a promising pathway to enhance agricultural sustainability by reducing input dependency and improving system efficiency. By identifying legume genotypes that promote beneficial downstream effects on cereals, breeders can select varieties that contribute residual nitrogen, improve soil structure, modify soil microbiome, and maintain moisture availability, thereby reducing resource inputs in subsequent crops. This approach can also extend beyond sequential rotations to intercropping systems where genetic variation in root architecture, nitrogen fixation efficiency, and rhizosphere interactions could enhance complementarity and resource-use efficiency. Whilst root traits have not been measured directly in this study, VOCs, soil properties and NDRE reflect substantial genotypic variability in processes likely mediated by root function. UAV-derived data and emerging technologies such as electromagnetic induction sensing and multispectral imaging have demonstrated potential to predict root functional traits and characterise root-zone properties at the population scale (Alahmad et al., 2025; Zhao et al., 2022), providing scalable phenotyping pathways for integrating root-mediated legacy effects into future breeding programs. The availability of modern genomic approaches now enable predictive breeding decisions (Hayes et al., 2024) that could harness legacy effects for the first time, and the ability to map these effects to defined genomic regions provides a foundation for incorporating them into genomic prediction pipelines.

However, implementing this approach at scale requires a paradigm shift that aligns breeding priorities across multiple crops or within large breeding programs that manage multiple crop portfolios simultaneously. Current breeding programs and funding models are structured around single-crop optimisation with economic incentives favouring direct trait improvements over legacy effects. Transitioning to legacy trait breeding would require new collaborative frameworks and revised intellectual property models that recognise contributions to farming systems productivity, alongside logistically complex multi-location, multi-season trials to validate genetic effect stability and build industry confidence. Legacy trait expression is likely to exhibit significant genotype-by-environment interactions, particularly when soil properties vary across target production environments, necessitating comprehensive testing to establish breeding value stability. This investment in comprehensive testing will be essential to build industry confidence and drive adoption of system-level breeding approaches.

As with any trait, studying and improving legacy effects requires sufficient genetic diversity. Elite breeding materials may lack the variation necessary to detect and exploit these effects compared to the mungbean diversity panel used in this research, necessitating the introgression of required genetic variation into breeding populations. This case study was also conducted at a single location with mungbean as the initial crop under nitrogen-deficient, rainfed conditions, which does not fully capture the complexity of legacy dynamics in diverse commercial farming systems. Nonetheless, yield from our mungbean mini-core experiments was similar to that of previous research experiments under different management conditions and years (Supplementary Fig 5). Furthermore, testing legacy effects across multiple wheat genetic backgrounds would reveal whether these effects are consistent across cereal genotypes or if specific legume-cereal genetic combinations maximise rotational benefits. Legacy effects will vary in their persistence across seasons as short-term effects mediated by residual mineral nitrogen may dissipate rapidly, whereas changes to soil microbiome structure or physical soil properties may endure across multiple growing seasons. Quantifying this temporal dimension will be important for breeding, as genotypes generating durable beneficial legacies would carry greater system-level value and warrant higher weighting within selection indices. Collectively, future studies involving multi-location trials, diverse cropping systems, and multiple species with appropriate genetic diversity are critical to explore the potential of this approach. Overall, this work provides the genetic and methodological foundation to enable breeding programs that optimise cropping systems rather than individual crops, offering a pathway for improved agricultural sustainability that harnesses genetic diversity to reduce external inputs while maintaining productivity.

## Methods

### Site preparation and nitrogen depletion

The experimental field site had characteristically high soil nitrogen levels typical of research stations due to fertiliser applications from previous trials. To create conditions more representative of commercial farming systems and conducive to evaluating genotypic variation in nitrogen fixation and legacy effects, soil nitrogen was deliberately depleted prior to sowing the mungbean panel. This was achieved by sequentially sowing grain sorghum and oat crops to maturity, which extracted residual soil nitrogen before being harvested and removed (Supplementary Fig 6). Soil sampling confirmed substantial nitrogen depletion before mungbean sowing (Supplementary Table 2), creating nitrogen-limiting conditions that would maximise expression of genotypic differences in nitrogen fixation capacity and enable quantification of legacy effects on the subsequent wheat crop.

### Plant material and experimental design

A diverse mungbean panel of 309 genotypes was sown on 22nd January 2024 at the Department of Primary Industries research station in Gatton, Australia (27.55 °S; 152.33 °E) (Supplementary Fig 7). This panel included the mungbean mini-core collection, a diverse subset from the World Vegetable Center genebank representing 19 countries across Asia, Australia, Africa and the Americas (Schafleitner et al., 2015). An additional 25 mungbean genotypes and one black gram (*Vigna mungo*) cultivar was also included (Supplementary Table 8). The trial used an unbalanced replicated design with the mini-core collection replicated twice and the additional genotypes replicated four times. The experiment was designed using a model-based row-column design that incorporated genetic relatedness among mungbean genotypes through a genomic relationship matrix derived from marker data. The design was optimised using the R package ’odw’ (Cullis et al., 2020) to account for genetic relatedness when allocating genotypes across the field.

Mungbean genotypes were sown in 2-row 1.52 m × 4 m plots (0.76m row spacing) at a density of 25 plants m□². Mungbean seeds were inoculated with Group I inoculum (EasyRhiz™, New Edge Microbials Pty. Ltd) at sowing to promote nitrogen fixation through rhizobia establishment in root nodules. Granulock Starter Z fertiliser (Incitec Pivot Fertilisers) was applied as a basal application of 25 kg ha□¹. The mungbean trial was desiccated with glyphosate and machine harvested to determine grain yield when the check cultivar Crystal reached 90% black pod stage. Harvest removed the majority of above-ground shoot biomass, with stubble retained in situ; no tillage was performed prior to wheat sowing, preserving the soil legacy conditions. Crystal was selected as the baseline mungbean genotype for its ‘typical’ phenology for the region aligning towards the centre of the commercial cultivars utilised in this study.

A single wheat cultivar (LongReach Raider) was sown 23 days after mungbean harvest (23rd May 2024) in the same paddock. No fertiliser was added to the wheat crop prior to, or during crop growth and maturity. Wheat plots (5-row, 1.52 m × 4 m with 0.3m row spacing) were positioned to exactly overlap with the previous mungbean plots, with a seeding rate of 120 plants m□². This spatial correspondence enabled traits measured in wheat to be directly mapped to the proceeding mungbean genotype. Upon maturity, wheat was machine harvested for grain yield determination. LongReach Raider was selected as it is a preferred cultivar for the northern growing region, where the experiment was conducted, being a widely grown Australian Prime Hard variety.

Both trials were conducted under rainfed conditions. Standard weed, insect and disease control was applied throughout both trials. No pest or disease outbreaks were observed. Daily meteorological data (minimum and maximum temperature, rainfall) were obtained from an onsite weather station (Supplementary Fig 1).

### Data Collection

Agronomic and physiological traits, as well as soil properties, were measured prior to and during both trials, either across the entire experiment or on a subset of diverse genotypes (Supplementary Fig 8).

#### Soil water and nutrient analyses

Prior to the commencement of the mungbean field trial, soil samples were taken from five soil cores across the paddock which were split into two depths (0-20 and 20-40cm) (Supplementary Table 9). Each sample was divided into two samples, the first was weighed, dried in 105°C oven, and then weighed again to calculate gravimetric water content (%). The second sample was used for analysis of extractable phosphorus (Colwell P), ammonium nitrogen (NH□□), and nitrate nitrogen (NO□□) using standard protocols described by Rayment and Lyons (2011). All analyses were performed by an accredited external laboratory using a SEAL AQ400 discrete analyser for colorimetric detection. All final concentrations were converted to mg/kg dry soil for reporting.

Colwell P was extracted using a 0.5 M sodium bicarbonate solution at a 1:50 soil-to-solution ratio. Samples were shaken for 16 hours, centrifuged, and filtered. Phosphorus concentrations in the extracts were determined colorimetrically using the ammonium molybdate–ascorbic acid method, with potassium antimonyl tartrate added to control the reaction rate. Concentrations measured in mg/L were converted to mg/kg based on extraction volume and soil mass.

Ammonium nitrogen (NH□□) was extracted using a 2 M potassium chloride (KCl) solution at a 1:10 soil-to-solution ratio and mixed for 1 hour. The resulting extracts were centrifuged and filtered before analysis. Ammonium was quantified colorimetrically through its reaction with sodium salicylate and nitroprusside in a mildly alkaline buffer, forming a coloured complex in the presence of chlorine and detected at 650 nm.

Nitrate nitrogen (NO□□) was extracted following the same procedure as ammonium using 2 M KCl. Nitrate was reduced to nitrite via a cadmium reduction column, and the resulting nitrite was measured colorimetrically using a reaction with sulfanilamide and NED, with absorbance read at 540 nm.

Additional soil nutrient analyses were undertaken prior to mungbean harvest from two replicates of 10 diverse mungbean genotypes (AGG324187 (2B), AusTRC 324159, Berken, Celera II-AU, CHIH-CO, CPI62672, Crystal, M10403, M10452 and Moong; Supplementary Fig 9). In addition to P, NH□□ and NO□□ being captured, exchangeable bases (Ca, K, Mg, Na) were also measured by using a 1 M ammonium chloride solution at a 1:10 soil-to-solution ratio, shaken for 1 hour. The extracts were centrifuged, filtered, and analysed using a Thermo iCAP PRO XP ICP-OES instrument. Concentrations (mg/L) were converted to mg/kg using extraction volume and soil mass. These values were further converted to milliequivalents (meq/100g) by dividing the mg/kg concentration by the atomic weight of each ion, and then adjusting for charge: divided by 10 for monovalent ions (Na□, K□) and by 5 for divalent ions (Ca²□, Mg²□). Cation exchange capacity (CEC) was estimated from the sum of these exchangeable cations.

#### Soil microbiome analyses

To investigate potential soil microbiome differences attributable to mungbean genotype, we analysed soil volatile organic compounds (VOCs) as chemical indicators of microbial activity and community composition. Soil samples were collected from the ten diverse genotypes described above, and VOC profiles were characterised using headspace solid-phase microextraction gas chromatography-mass spectrometry (HS-SPME-GC/MS).

The HS-SPME-GC/MS methodology was adapted from Rivers et al. (2019) for volatilomic analysis of soil samples. Analysis was performed using an Orbitrap Exploris GC 240 Mass Spectrometer fitted with a TriPlus RSH Multipurpose (i.e. Liquid and Headspace Injection, SPME Arrow, and ITEX-DHS) Extended Rail (Thermo Fisher Scientific, Massachusetts, USA). Splitless mode was employed for application of the VOCs via a 1.0 mm straight no-wool liner to a TRACE 1610 series Gas Chromatograph (Thermo Fisher Scientific). Soil samples, QC samples, laboratory blanks, and standards were block randomised throughout each batch. Each sample was placed on 2 ml of 20% NaCl solution and incubated at 80 °C for 40 minutes with continuous agitation at 500 rpm to enhance VOC volatilisation and homogenisation. VOC extraction (adsorption) was carried out by exposing a broad specificity SPME Arrow fibre (1.10mm DVB/C-WR/PDMS 110μm x 20mm, Thermo Fisher Scientific) to soil sample head space for 40 minutes with continuous agitation at 1000 rpm. Extracted VOCs underwent desorption from the SPME Arrow on to the GC column (TraceGOLD TG-5SilMS GC Column, Thermo Fisher Scientific) via the inlet at 250 °C for 10 minutes. Between samples, the SPME Arrow fibre undergoes a pre- and post-conditioning at 250 °C for an additional 10 minutes to prevent cross sample contamination. Chromatographic separation of the soil VOCs was achieved employing ultra-pure helium as a carrier gas and via a 38.5 minutes oven temperature program: 40 °C for 2 minutes, increasing to 150 °C at a ramp rate of 5 °C/min and held for 2 minutes, then increased to 320 °C at 15 °C/min and held for 1 minute. Lastly, the Orbitrap Exploris 240 MS was operated in full MS scan acquisition using electron ionisation at 70eV from m/z 40-500 with a lag time of 5 minutes, and a Orbitrap resolution of 60,000. Temperatures were 320 °C for the auxiliary transfer line and 250 °C for the ion source.

Volatile data were processed using Thermo Compound Discoverer (ver. 3.3.2.31) for deconvolution and data analysis. The NIST/EPA/NIH Mass spectral libraries (ver. 2023) were used for mass spectral matching and peak annotations. Kovats non-isothermal retention indices (RIs) were calculated using an *n*-alkanes (C7-C40) standard mix and compared to reference semi-standard non-polar RIs from the NIST and PubChem databases when available. Peak intensity values were collected and corrected based on the internal standard nitrobenzene, background peak detection in blank samples, and sample weight. Subsequent statistical analyses were carried out using MetaboAnalyst 6.0 (www.metaboanalyst.ca).

#### Phenology

Days to flowering was captured for all plots within the mungbean trial and was determined when 50% of the plants within the plot had flowered (Dudley et al. 2025). For the subsequent wheat crop, time to flowering was determined according to GS65 of the Zadoks scale (Zadoks et al. 1974).

#### Total aboveground biomass

In the mungbean trial, biomass samples were harvested from 1-4 replications of a subset of 47 lines, representing a genetically diverse set of germplasm and breeding lines previously used in physiology studies, including the reference cultivar Crystal. Sampling occurred when the commercial check cultivar Crystal reached flowering (50% of the plot had open flower), which represented mid canopy development. Biomass was collected from a 0.5 m section at the end of the plot spanning both rows (0.5 × 1.52 m). For the wheat trial, biomass cuts were taken from the same plots as the preceding mungbean trial at heading. Samples were collected from a 0.5 m section at the end of the plot from the two middle rows, corresponding to a biomass sampling area of approximately 0.5 × 0.75 m. For both mungbean and wheat biomass samples, they were dried at 65 °C for 5 days before weighing total biomass.

#### Vegetation indices

Vegetation indices were extracted from multispectral UAV imagery collected when mungbean check cultivar Crystal reached flowering and 90% black pod growth stages. Imagery collection and processing followed methods detailed in Van Haeften et al. (2025). Briefly, a MicaSense Altum™ sensor mounted on a Matrice 300 RTK UAV platform captured images that were processed into georeferenced orthomosaics using Pix4D software. Plot boundaries were defined in ArcMap for the mungbean trial and applied to the wheat trial to ensure corresponding wheat plots aligned with previous mungbean genotype plots. Normalized difference red edge index (NDRE), height, thermal and canopy coverage were extracted and used to estimate mungbean biomass at mid-canopy development (determined when Crystal had reached 50% flowering). Raw NDRE values were also extracted at 90% black pod stage in mungbean and at flowering in wheat. Mungbean NDRE was measured at this stage as a proxy for end of season resource allocation. NDRE was calculated as NDRE= (NIR -RedEdge)/(NIR+RedEdge), where NIR is Near Infrared Reflectance and RedEdge is the red edge spectral band.

#### Grain quality characteristics

To assess legacy effects on wheat quality in addition to yield, grain protein content was measured. Wheat grain protein content was determined using near-infrared spectroscopy with an Inframatic 9500 (IM9500) Grain Analyser (Perten Instruments), a commercially validated instrument widely used in Australian grain quality assessment. The instrument warmed up for a minimum of 30 minutes and self-calibrated using the internal reference standard per manufacturer recommendations. Approximately 200 g sub-samples were loaded into the IM9500 sample hopper for analysis The IM9500 conducted three sub-sample readings and averaged spectra before predicting constituent values. Each sample was analysed in triplicate; between replicates, the hopper was emptied and refilled with fresh sample portions. Protein content was reported on a 12% moisture basis.

## Statistical Analyses

### Mungbean biomass prediction models

Mungbean biomass for all plots was predicted using the multispectral data captured for the UAV flight corresponding to mid-canopy development and calibrated against the ground-based biomass measurements collected from the genetically diverse 47 genotypes on the same day. A prediction model previously developed for this growth stage (Van Haeften et al., 2025) was applied, incorporating height, thermal, NDRE, and coverage in a stepwise regression framework. This approach enables estimation of biomass as a more direct indicator of plant physiological performance than vegetation indices alone. Model validation used an 80/20 training/testing split to determine the accuracy of the model in predicting biomass for this trial

### Single-site spatial analyses

Single site spatial analysis accounted for spatial variation across the site for all traits in both crops. Both fixed and random row and column terms were tested for model improvement and retained as needed. Wald statistics determined significance of fixed effects, while AIC, BIC and log-likelihood assessed model fit changes due to added random components. Best Linear Unbiased Estimators (BLUEs) with genotype as a fixed effect and Best Linear Unbiased Predictors (BLUPs) with genotype as a random effect were calculated using equation 1:

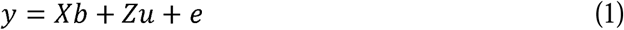

where b is a vector of fixed effects, u is a vector of random effects, X and Z are the associated design matrices and e is the residual.

Generalised broad sense heritability for each trait was calculated following Cullis et al (2006) using equation 2:

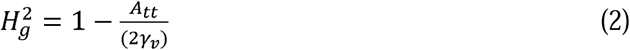

where A_tt_ is the average prediction error variance and r_V_ is the genetic variance estimated by the model.

### Correlations

Phenotypic correlations between traits were calculated using Pearson’s correlation coefficient based on BLUEs for each genotype. Statistical significance was assessed using two-tailed p-values, with non-significant correlations (p > 0.05) flagged in visualisations. The same methodology was used to calculate correlations between haplotype variances and haplotype effects.

Genetic correlations were calculated for each trait pair using mixed linear models with unstructured variance structures applied to trait-to-trait interaction terms as per equation 3:

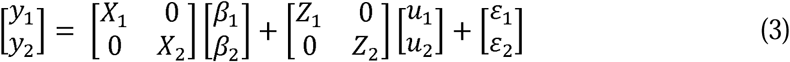

where y represents phenotype vectors for traits 1 and 2 respectively, X and Z are design matrices for the fixed and random components of the model, β represents coefficient of fixed effects, u represents the random genetic effects, and *_E_*, is the residual. Covariance between genetic effects is represented as var(u) =G ⊗ K, where K is the genomic relationship matrix (GRM) and G is the genetic covariance matrix between traits.

### Genomic analyses

To identify genomic regions underlying both mungbean performance and legacy effects on wheat, haplotype-based mapping was conducted using the local genomic estimated breeding value (local GEBV) method (Voss-Fels et al., 2019; Shaffer et al., 2025). DArTseq markers were curated based on a minor allele frequency greater than 0.05, heterozygosity of less than 0.2, and <20% missing calls, resulting in 5,516 high-quality markers. Linkage disequilibrium (LD) was explored to identify suitable thresholds for haploblock definition. The R package ‘Selection Tools’ defined haploblocks with markers in LD >0.3 and a marker tolerance of 4. These parameters were chosen due to low levels of LD in the population, given the diverse nature of mungbean genotypes used in the study and to ensure blocks remained a useful size for implication in practical breeding contexts. Marker tolerance represents the maximum number of markers that can be skipped during haploblock creation, chosen based on the potential for misalignment. This approach defined 1,126 haploblocks with an average of 4.9 markers per block.

Marker effects were calculated using ridge-regression BLUP to determine simultaneous effects of all SNPs per equation 4.

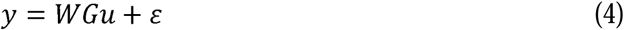

where u is a vector of marker effects, G is the genotype matrix, W is a design matrix relating genotypes to observations (y), and ε is the error. Relevant SNP effects were summed to calculate haplotype effects at each haploblock. Effect variance was calculated for each haploblock. For each trait, the top 1% of haploblocks by variance (12 haploblocks) were identified as the most important.

### Identifying and co-locating mungbean orthologues for functional legume genes

Minimap2 (Li, 2018) was used to extract regions within high variance haploblocks based on the Crystal reference genome enhanced with additional PacBio HiFi long-read sequencing data (C. Mens & M. Udvardi, unpublished data). Mungbean orthologues of known nodulation and nitrogen fixation genes were identified using reciprocal tBLASTn searches (E-value threshold ≤ 1e-5) against a compiled list of over 200 characterised genes in model legume systems (Roy et al., 2022). Orthologues were retained if overlapping with at least one of five key haploblocks selected for their high variance for wheat yield and/or protein content.

### Weighted selection strategies simulations

Three breeding program scenarios were simulated to evaluate genetic improvement strategies: 1) selection for mungbean yield alone, 2) selection for legacy wheat yield alone, and 3) simultaneous improvement of both crops. For the simultaneous improvement scenario, selection used trait indices calculated as weighted sums of breeding values with mungbean:wheat weightings of 0.7:0.3, 0.5:0.5, and 0.3:0.7. Simulations used SNP effects estimated from genomic prediction models to calculate genomic estimated breeding values (GEBVs). Each program ran 25 cycles, selecting 30 parents per cycle based on program-specific criteria. Selected parents were randomly mated to create 50 crosses, with 100 progeny per cross, generating 5,000 selection candidates for the next cycle. Genetic improvement was assessed by population mean GEBV changes per cycle. Each program was simulated five times, reporting average values and 95% confidence intervals of population mean GEBVs using the genomicSimulation R package (Villiers et al., 2022).

### Comparison of genotypic differences for soil characteristics

To evaluate whether soil characteristics differed significantly among the subset of 10 diverse mungbean genotypes described in the soil nutrient analyses section, one-way analysis of variance (ANOVA) was performed for each measured soil variable. Genotype was treated as a fixed effect, and significance was assessed to determine whether plant genetic variation contributed to differences in soil nutrient concentrations and moisture content.

## Acknowledgments

We thank the World Vegetable Center (WorldVeg), particularly Roland Schafleitner and Ram Nair, for providing the genomic data for the mungbean mini-core collection that formed the foundation of this study. We are grateful to the Queensland Department of Primary Industries for Gatton field trial support and provision of the Hermitage 2018 yield datasets. We particularly acknowledge the technical staff at the Gatton Research Facility for their assistance with field operations, and trial management. We thank LongReach Plant Breeders for generating the wheat grain protein data.

## Data availability

All primary data supporting the findings of this study are available via The University of Queensland eSpace repository at https://doi.org/10.48610/cee8b66.

## Funding

This work was supported by The University of Queensland start-up funds awarded to MS. Mungbean research conducted by MS, LH, MU at The University of Queensland was supported by the Australian Centre for International Agricultural Research International Mungbean Improvement Network Phase 2 (CIM2014/079) and Grains Research & Development Corporation Genetic Initiative to Transform Symbiotic Nitrogen Fixation in Australian Pulse Crops (UOQ2403-012RTX). LH was supported through an ARC Future Fellowship (FT220100350). ED was supported by the Queensland Government’s Industry Research Fellowships program (AQIRF096-2023RD6). SVH, ME were recipients of Grains Research & Development Corporation Research Scholarships (UOQ2101-003RSX; UOQ2410-011RSX). SVH, SMB were recipients of an Australian Government Research Training Program Scholarship. SMB was supported as a PhD student by the Deutsche Forschungsgemeinschaft (DFG) and University of Queensland International Research Training Group 2843 – Accelerating Crop Genetic Gain. ME was supported as a PhD student of the Australian Research Council Training Centre in Predictive Breeding for Agricultural Futures (IC230100016).

## Authors contributions

**Shanice Van Haeften:** Writing – Original Draft Preparation, Investigation, Data Curation, Formal analyses, Visualisation. **Stephanie M. Brunner:** Writing – Original Draft Preparation, Data Curation, Formal analyses, Visualisation. **Eric Dingalsan:** Methodology, Writing – Review & Editing. **Edward Fabreag:** Formal analyses, Writing – Review & Editing. **Joseph Eyre:** Methodology, Investigation, Writing – Review & Editing. **Celine Mens:** Investigation**. Ben J. Hayes:** Supervision, Writing – Review & Editing. **Michael Udvardi:** Supervision, Writing – Review & Editing. **Samir Alahmad:** Investigation. **Mitchell Eglinton:** Investigation, Writing – Review & Editing. **Ryan McQuinn:** Investigation, Formal analyses, Writing – Review & Editing. **Merril Ryan**: Resources, Writing – Review & Editing. **Sarah van der Meer:** Investigation. **Millicent R. Smith:** Conceptualisation, Funding Acquisition, Project Administration, Writing – Review & Editing. **Lee T. Hickey:** Conceptualisation, Project Administration, Writing – Review & Editing.

## Competing interests

The authors declare no competing interests.

## Notes

https://doi.org/10.48610/cee8b66

### Competing Interest Statement

The authors have declared no competing interest.

### Summary of Updates

Revised version incorporating feedback from external reviewers.

