## Supplementary Figures for "The potential to breed for genetic legacy effects in sustainable farming systems"

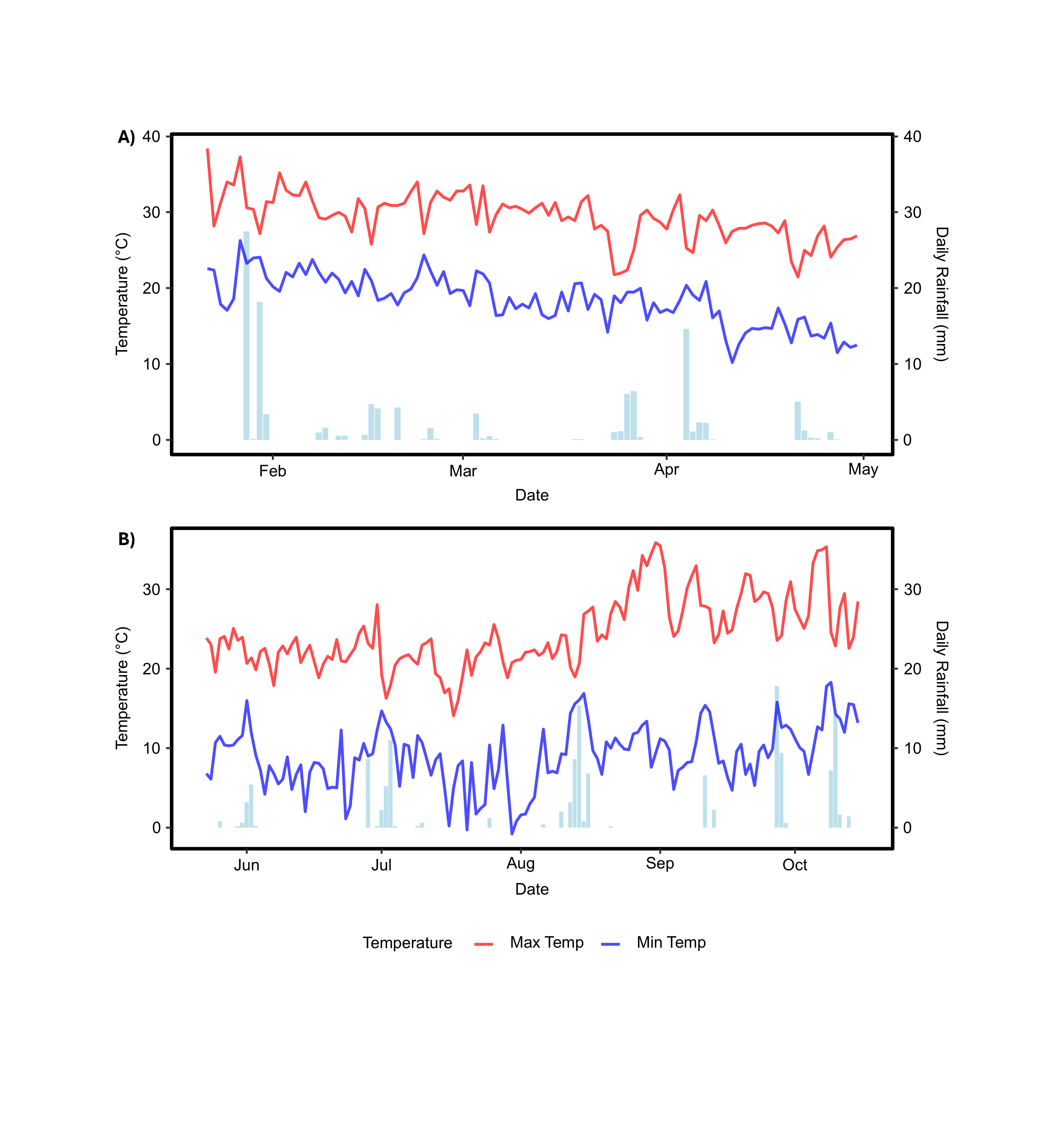

**Supplementary Fig 1.** Daily maximum temperature (red, °C), minimum temperature (blue, °C) and rainfall (bar charts, mm) data from sowing to harvest date for mungbean (A) and wheat (B) trials at Gatton, Queensland, Australia.

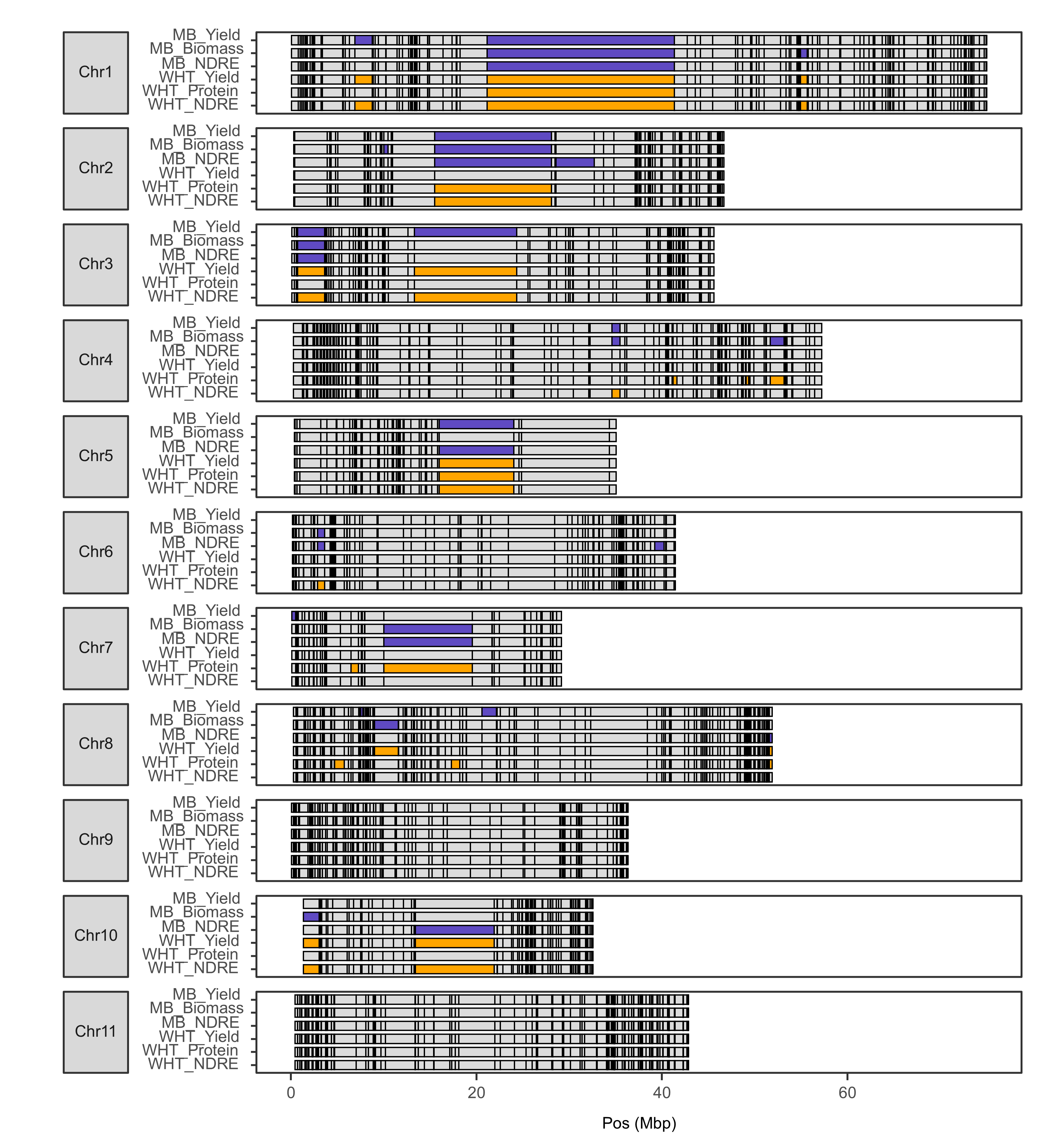

**Supplementary Fig 2.** Genomic regions associated with additional physiological and legacy traits in mungbean and wheat. The top 1% of high-variance haploblocks are highlighted in purple for mungbean traits and in orange for wheat traits.

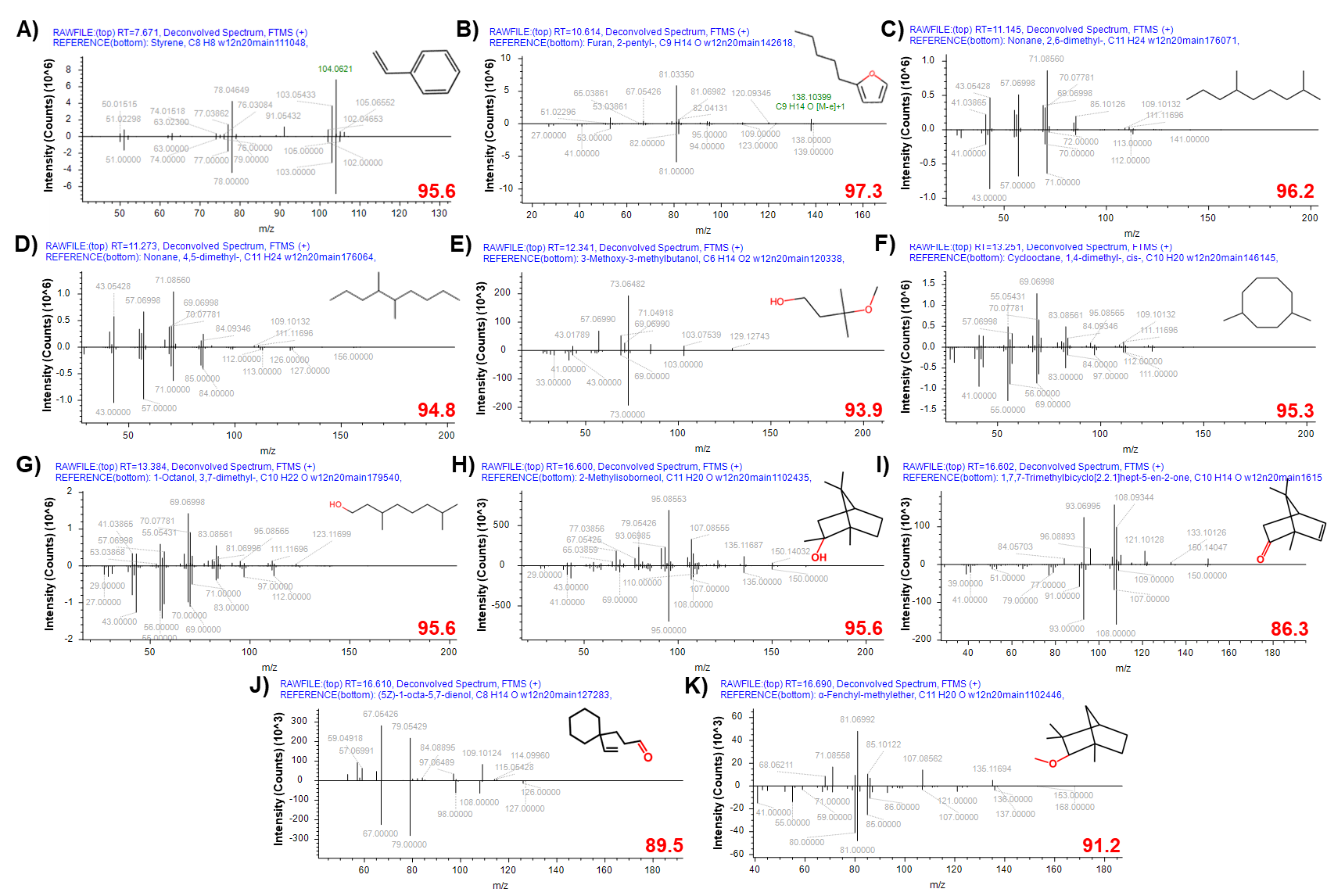

**Supplementary Fig 3.**  MS fingerprint and NIST match of the soil VOCs displaying statistical significance between the 11 genotypes tested and the Bare Soil control (A) styrene; (B) furan, 2-pentyl; (C) nonane, 2,6-dimethyl; (D) nonane, 4,5-dimethyl; (E) 3-methoxy-3-methylbutanol; (F) cyclooctane, 1,4-cis-dimethyl; (G) 1-octanol, 3,7-dimethyl; (H) 2-methylisoborneol; (I) dehydrocamphor; (J) (5Z)-1-octa-5,7-dienol (K) a-fenchyl-methylester. Compound structures can found in the top right corner of the MS plot, and in red at the lower right corner is the total score (0-100, combines several metrics including: search index, reverse search index, high resolution filtering and reverse high resolution filtering) for each NIST match.

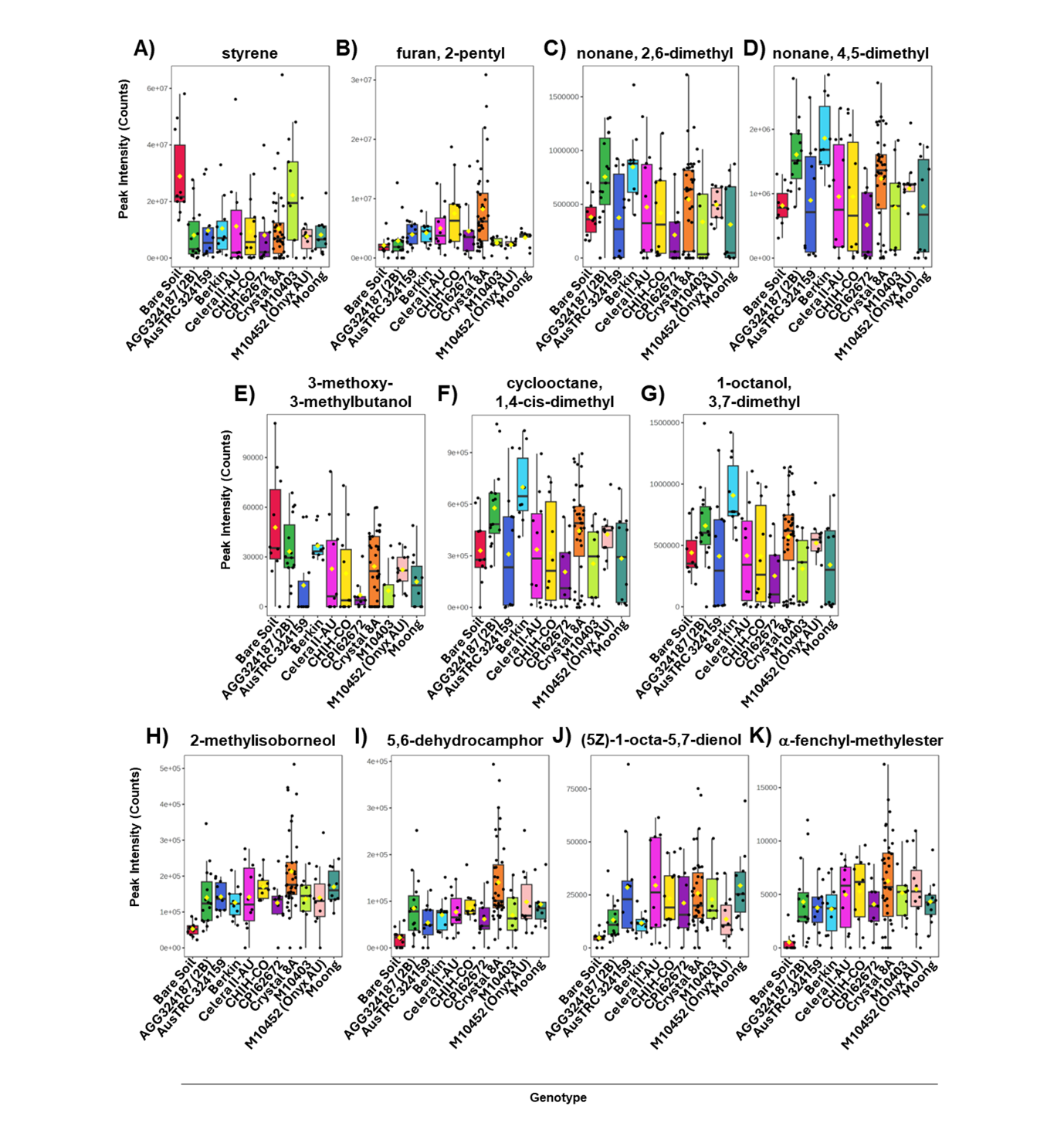

**Supplementary Fig 4.** Differential peak intensities of soil VOCs displaying statistical significance between the subset of 11 genotypes analysed and the Bare Soil control, (A) styrene; (B) furan, 2-pentyl; (C) nonane, 2,6-dimethyl; (D) nonane, 4,5-dimethyl;( E) 3-methoxy-3-methylbutanol; (F) cyclooctane, 1,4-cis-dimethyl; (G) 1-octanol, 3,7-dimethyl; (H) 2-methylisoborneol; (I) dehydrocamphor; (J) (5Z)-1-octa-5,7-dienol (K) a-fenchyl-methylester.

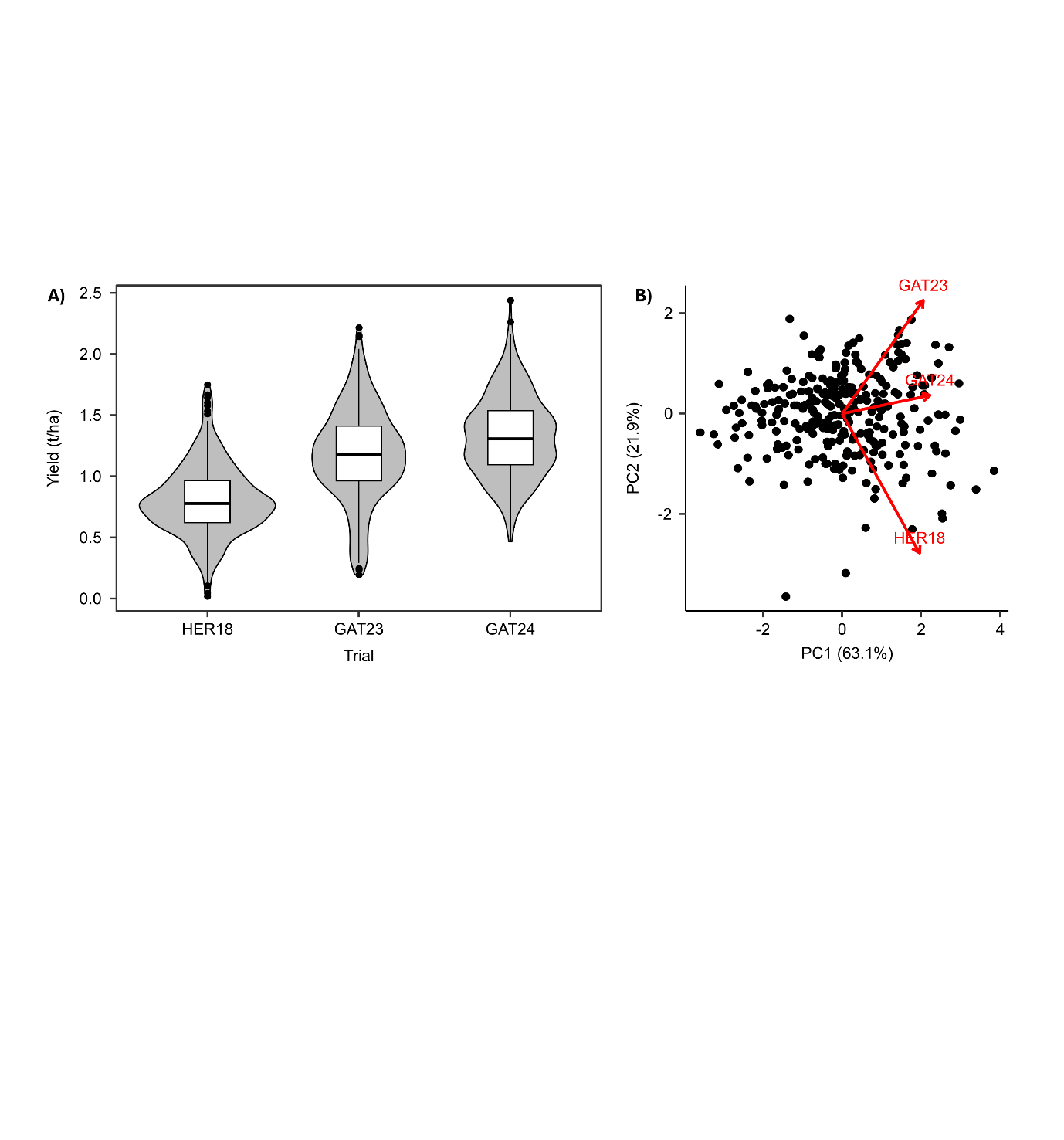

**Supplementary Fig 5.** Comparison of mungbean yield across experiments and principal component analysis (PCA) of site variation. (A) Boxplots of historical yield data (BLUEs) from the current mungbean field trial alongside two previously conducted field trials using the same mini-core panel under non-nitrogen-limiting conditions. This comparison demonstrates that yield performance in the current study is consistent with historical trials, confirming the representativeness of the experimental conditions. (B) PCA plot based on yield BLUEs across all sites.

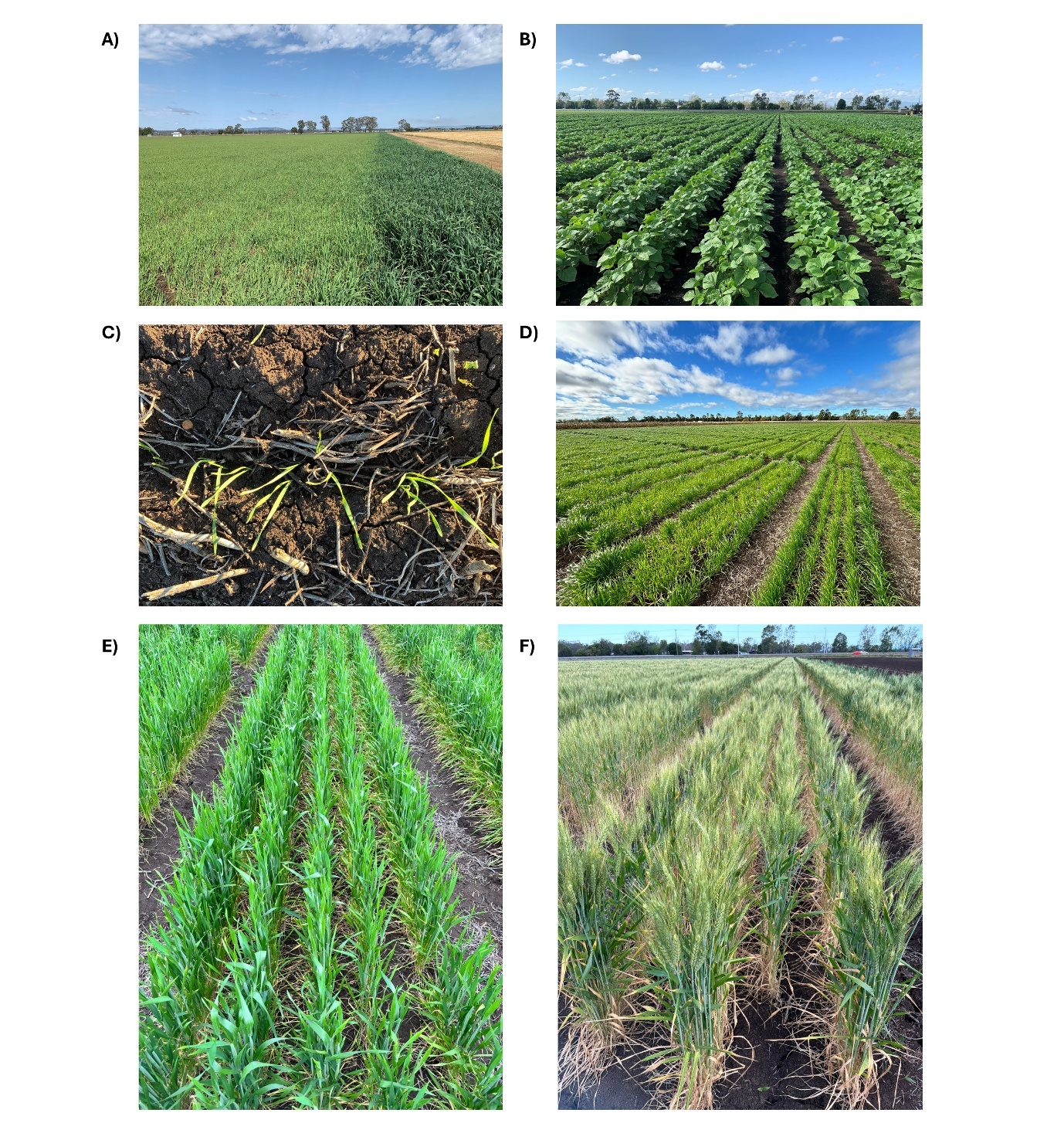

**Supplementary Fig 6.** Photographs illustrating field trial conditions and crop growth stages across the mungbean-wheat rotation experiment.**.** (A) Visual contrast between the nitrogen-depleted trial area following the oat cover crop (left) and the adjacent area without cover cropping (right), highlighting boundary effect, (B) Mungbean yield trial at peak flowering stage, (C) Emergence of wheat seedlings from mungbean stubble, illustrating early establishment conditions, (D) Wheat trial during early vegetative growth , (E) Representative wheat plot at booting stage , (F) Representative wheat plot at heading stage.

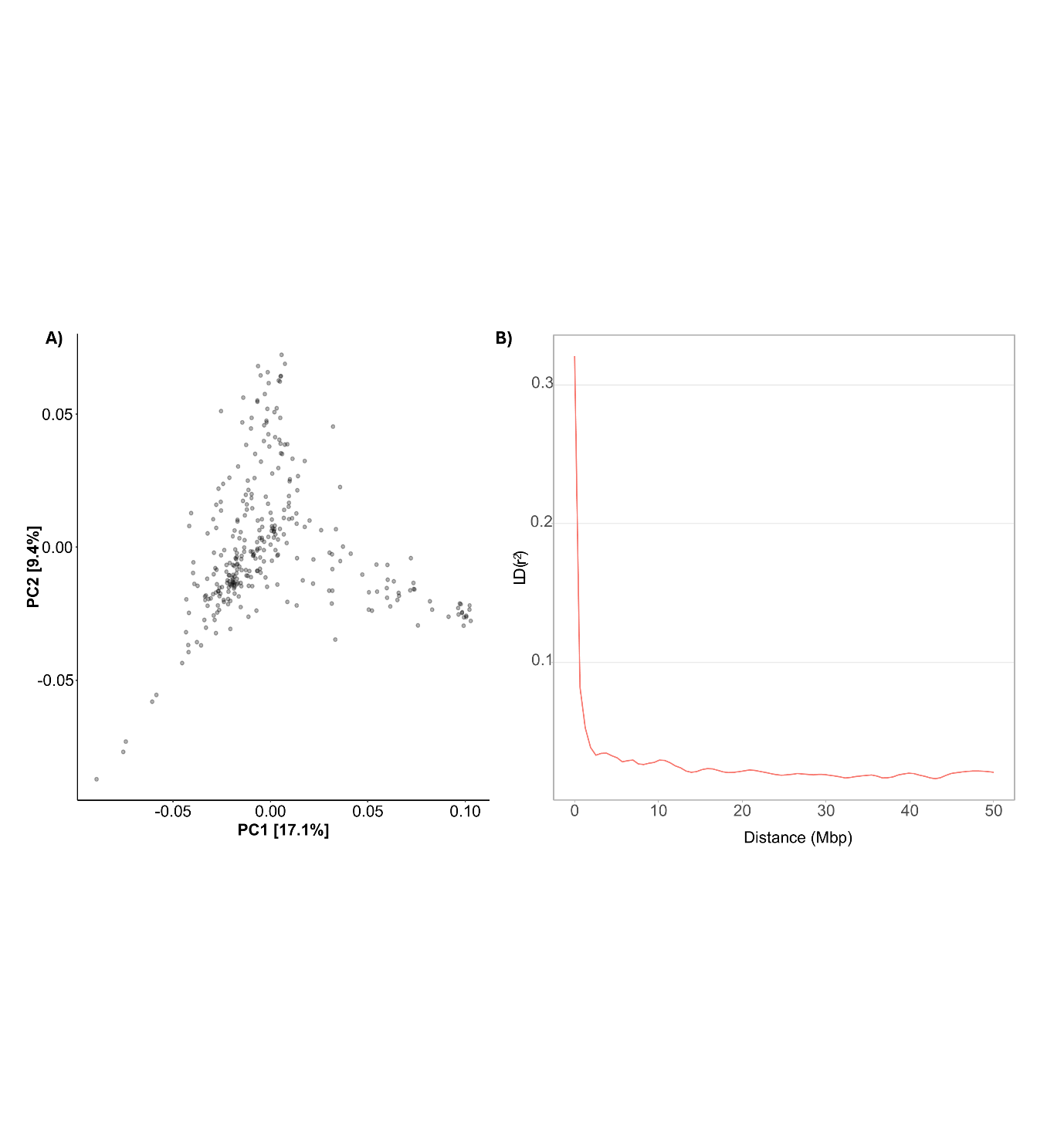

**Supplementary Fig 7.** Population structure and linkage disequilibrium decay of the mungbean mini-core set based on genome-wide SNP data, showing (A) a principal component analysis (PCA) illustrating genetic differentiation among genotypes used in the study and (B) mean r² values for SNP pairs within 100 kb.

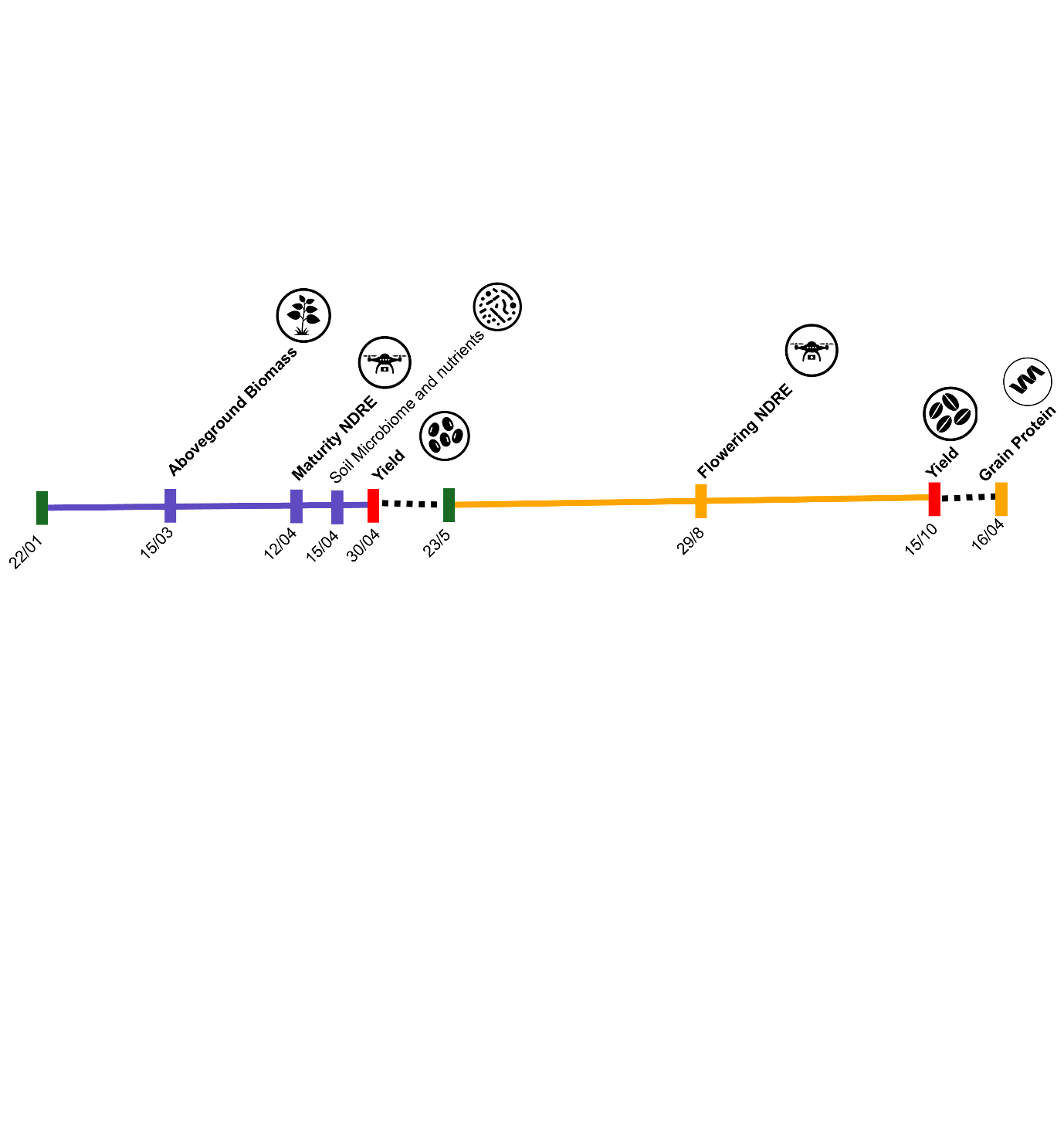

**Supplementary Fig 8.** Timeline of physiological and UAV data collection captured across mungbean and wheat trials.

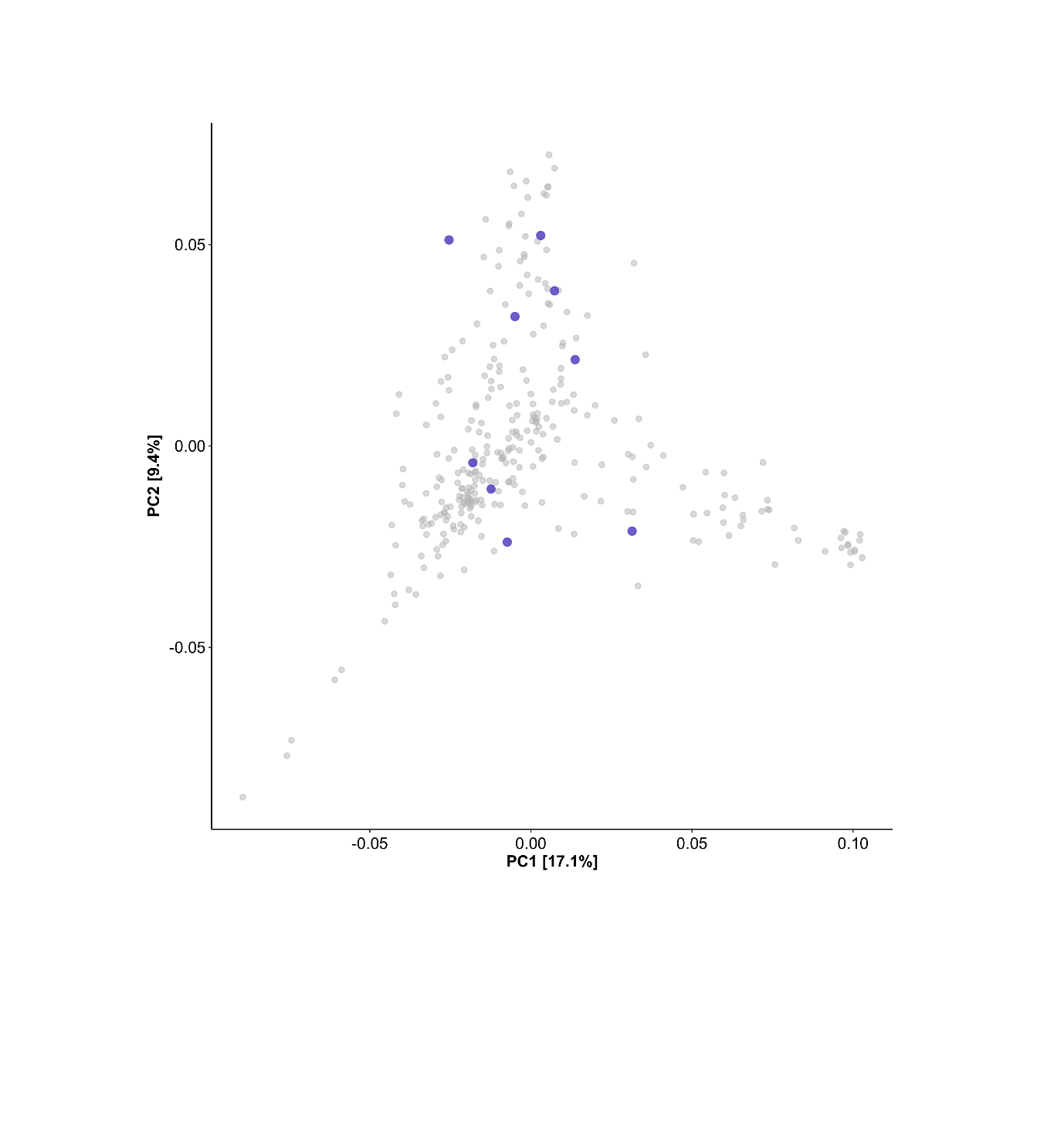

**Supplementary Fig 9.** **Principal Component Analysis (PCA) of genome-wide diversity among mungbean genotypes based on SNP markers**. The nine selected genotypes used for detailed soil nutrient and microbiome analysis are highlighted in blue, positioned within the broader mungbean diversity panel (grey). One additional selected genotype (Onyx) was excluded from the plot due to unavailable marker data.
